# Crystal structures of bacterial Small Multidrug Resistance transporter EmrE in complex with structurally diverse substrates

**DOI:** 10.1101/2022.01.11.475788

**Authors:** Ali A. Kermani, Olive E. Burata, B. Ben Koff, Akiko Koide, Shohei Koide, Randy B. Stockbridge

**Author notes:** Equal contributions (Ali A. Kermani, Olive E. Burata).

## Abstract

Proteins from the bacterial small multidrug resistance (SMR) family are proton-coupled exporters of diverse antiseptics and antimicrobials, including polyaromatic cations and quaternary ammonium compounds. The transport mechanism of the *Escherichia coli* transporter, EmrE, has been studied extensively, but a lack of high-resolution structural information has impeded a structural description of its molecular mechanism. Here we apply a novel approach, multipurpose crystallization chaperones, to solve several structures of EmrE, including a 2.9 Å structure at low pH without substrate. We report five additional structures in complex with structurally diverse transported substrates, including quaternary phosphonium, quaternary ammonium, and planar polyaromatic compounds. These structures show that binding site tryptophan and glutamate residues adopt different rotamers to conform to disparate structures without requiring major rearrangements of the backbone structure. Structural and functional comparison to Gdx-Clo, an SMR protein that transports a much narrower spectrum of substrates, suggests that in EmrE, a relatively sparse hydrogen bond network among binding site residues permits increased sidechain flexibility.

## Introduction

The small multidrug resistance (SMR) family of microbial membrane proteins is a well-studied family composed of primitive dual-topology proton-coupled transporters. The SMR family has two major physiological subtypes that can be distinguished based on sequence(Kermani et al., 2020). Representatives of the “Gdx” (<u>g</u>uani<u>d</u>inium e<u>x</u>port) subtype export a bacterial metabolite, guanidinium ion (Gdm^+^), in exchange for two protons(Kermani et al., 2018; Nelson et al., 2017). Representatives of the “Qac” (<u>q</u>uaternary <u>a</u>mmonium <u>c</u>ompound) subtype are proton-coupled exchangers of quaternary ammoniums and other hydrophobic, cationic compounds. Since the first quaternary ammonium antiseptics were introduced approximately one hundred years ago, proteins from the Qac cluster have been closely associated with the spread of multidrug resistance elements (Gillings, 2017; Pal et al., 2015; Russell, 2002; Zhu et al., 2017).

Many bacteria possess SMR proteins belonging to both subtypes. Transporters from the Qac and Gdx clusters do not overlap in terms of physiological role: the Qac proteins do not transport Gdm^+^ and require additional hydrophobicity in transported substrates, whereas the Gdx transporters require substrates to have a guanidinyl moiety and cannot export quaternary ammoniums or other cations(Kermani et al., 2020). However, the two subtypes transport an overlapping subset of hydrophobic substituted guanidinium ions and share high sequence conservation (∼35% sequence identity), strongly suggesting conservation of the overall fold.

The best-studied of the Qac proteins is the *E. coli* transporter, EmrE. The substrate repertoire of EmrE includes planar, conjugated aromatic ring systems, quaternary ammoniums and phosphoniums (with or without aromatic substituents), and substituted guanidiniums. EmrE also provides resistance to biocides from these substrate classes with long alkyl tails, such as benzalkonium and cetyltrimethylammonium, which are found in common household antiseptics. Although mechanisms to explain the transport promiscuity have been proposed, (Jurasz et al., 2021; Robinson et al., 2017), the structural basis for substrate binding is unknown, and for many years, structural information was limited to low-resolution models without loops or sidechains(Fleishman et al., 2006; Ubarretxena-Belandia et al., 2003), impeding a full description of the molecular mechanism. A previous crystal structure of EmrE was unreliable for molecular analysis, with no sidechains modelled, poor helical geometry, and helices too short to span the membrane (Chen et al., 2007). Computational models constrained by the low-resolution data have also been proposed (Ovchinnikov et al., 2018; Vermaas et al., 2018). Recently, high resolution structural information for the SMR family has begun to emerge. First, crystal structures of a Gdx homologue from *Clostridales*, Gdx-Clo, were resolved in complex with substituted guanidinium compounds including octylguanidinium (Kermani et al., 2020). In addition to revealing the binding mode of the guanidinyl headgroup, the structure of Gdx-Clo with octylguanidinium showed that hydrophobic repacking of residues lining one side of the binding pocket opens a portal from the substrate binding site to the membrane interior, accommodating the substrate’s long alkyl tail. In addition, a model of an EmrE mutant with reduced conformational exchange dynamics, S64V, computed from extensive NMR measurements, was also reported recently (Shcherbakov et al., 2021).

Here we report several crystal structures of EmrE, including a low-pH (proton-bound) structure and five structures in complex with structurally diverse quaternary phosphonium, quaternary ammonium, and planar aromatic substrates. Structure determination was facilitated by repurposing a monobody crystallization chaperone that we originally developed for Gdx-Clo(Kermani et al., 2020). The various substrates are accommodated by EmrE with minimal changes in the backbone structure. Instead, binding site tryptophan and glutamate sidechains adopt different rotamers to accommodate different drugs. These sidechain motions expand or reduce the binding pocket and provide ring-stacking interactions for structurally disparate substrates. We propose that, compared with the closely related but more selective SMR transporter, Gdx-Clo, a reduced network of hydrogen bond interactions in the EmrE binding site allows sidechain flexibility to accommodate substrates of different shapes and sizes without requiring substantial alteration of EmrE’s backbone configuration.

## Results

### Engineering of EmrE to introduce a monobody binding site

We recently solved a crystal structure of a metabolic Gdm^+^ exporter from the SMR family, Gdx-Clo (Kermani et al., 2020). For this effort, we selected monobody crystallization chaperones from large combinatorial libraries (Koide et al., 2012; Sha et al., 2017), which aided in crystallization of the transporter. Upon structure determination, we noticed that the epitope of Gdx-Clo recognized by monobody L10 is limited to a 9-residue stretch of loop 1 that is relatively well-conserved among SMR proteins (Figure 1A). Moreover, crystal contacts are mediated almost entirely by the monobody, whereas contacts between the transporter and a symmetry mate are limited to just five hydrophobic residues contributed by TM4_A_ and TM4_B_ (Figure 1 – Figure Supplement 1). These observations suggested that conservative mutagenesis of EmrE loop 1 to introduce the Gdx-Clo residues might permit monobody L10 binding in order to facilitate crystallization of EmrE. We therefore designed a triple mutant, E25N, W31I, V34M, which we call EmrE_3_. Previous studies showed minimal functional perturbation upon mutation of E25 and W31 to Ala or Cys(Elbaz et al., 2005; Yerushalmi and Schuldiner, 2000). All three residues are predicted to be a loop distant from the substrate binding site, and none of the three are conserved in the SMR family.

**Figure 1.**
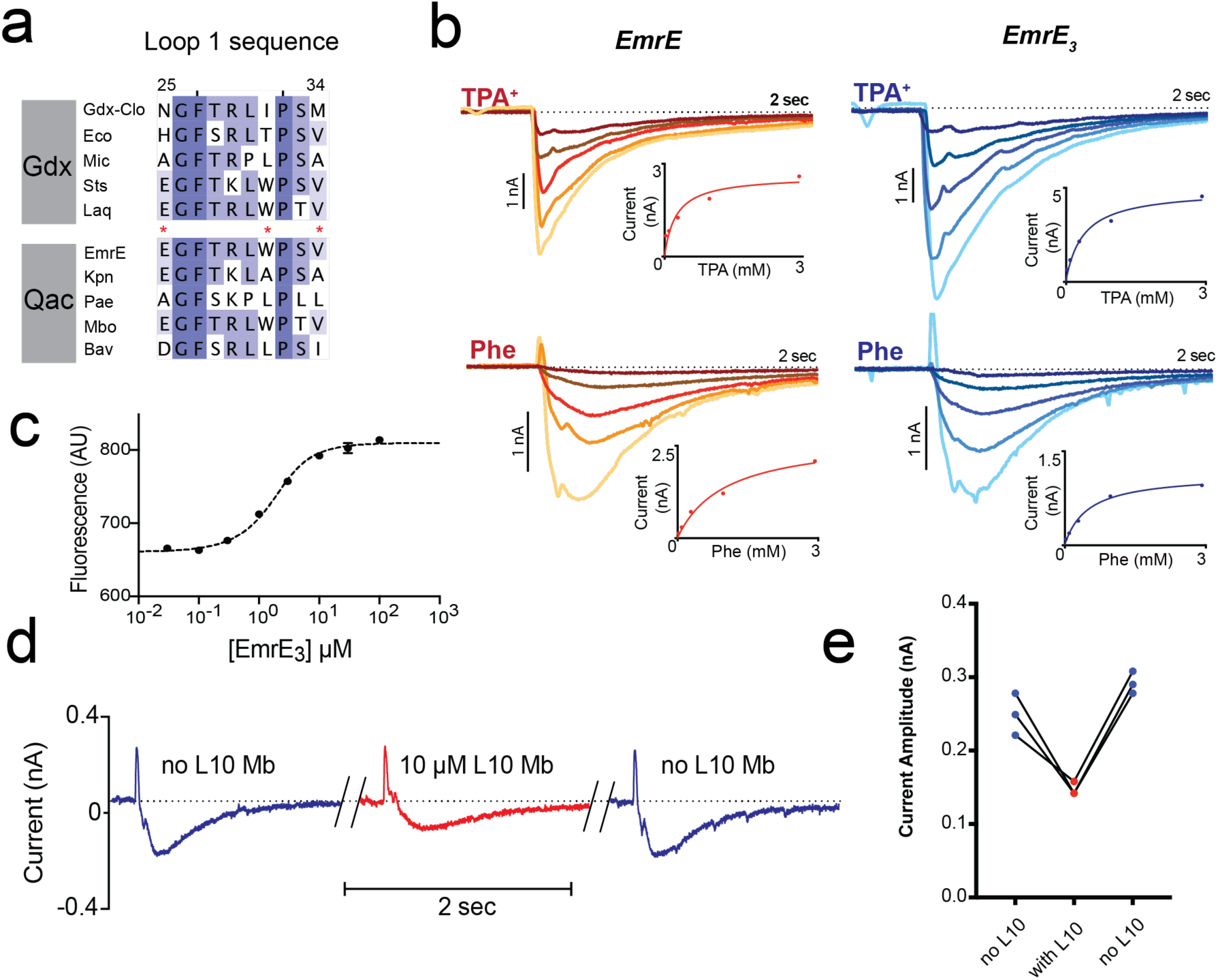
Introduction of monobody binding epitope to EmrE. A. Sequence alignment for loop 1 of selected SMR proteins, numbered according to EmrE sequence. From top to bottom: representative Gdx sequences (*Clostridiales* bacterium oral taxon 876, *Escherichia coli*, *Micromonospora*, *Streptomyces tsukubensis*, and *Leifsonia aquatica*) and representative Qac sequences (*Escherichia coli*, *Klebsiella pneumoniae*, *Pseudomonas aeruginosa*, *Mycobacterium bovis*, and *Bordetella avium*). Positions mutated in the EmrE_3_ construct (E25N, W31I, V34M) are indicated with red asterisks. B. Representative currents evoked by perfusion of WT EmrE or EmrE_3_ sensors (shades of red and blue, respectively) with 30 μM – 3 mM TPA^+^ (top panels) or PheGdm^+^ (Phe, lower panels). Insets show plot of peak current amplitude as a function of substrate concentration for a representative titration performed using a single sensor. Solid lines represent fit of datapoints from a single titration series to the Michaelis-Menten equation. K_m_ values for independent sensor replicates are reported in Figure 1-Figure Supplement 2. C. Microscale thermophoresis measurement of EmrE_3_ binding to monobody L10. Points and error bars represent mean and SEM of three independently prepared samples. Where not visible, error bars are smaller than the diameter of the point. Dashed line represents fit to Eq. 1 with K_d_ = 850 nM. Representative raw data trace is shown in Figure 1 – Figure Supplement 3. D. EmrE_3_ currents evoked by 1 mM PheGdm^+^. Sensors were incubated for 10 minutes in the presence (red traces) or absence (blue traces) 10 μM monobody L10 prior to initiating transport by perfusion with PheGdm^+^. Currents shown are from a representative experimental series using a single sensor preparation. E. Peak currents measured for three independent perfusion series performed as in panel D. Peak currents decreased an average of 40 ± 1.5 % in the presence of monobody.

In accord with these observations, solid supported membrane (SSM) electrophysiology experiments showed that the EmrE_3_ variant is active and transports representative substrates tetrapropylammonium (TPA^+^) and phenylguanidinium (PheGdm^+^). Upon perfusion with substrate, negative capacitive currents are evoked, indicating an electrogenic transport cycle, with substrate transport coupled to the antiport of ∼2 H^+^, as has been previously reported for these(Kermani et al., 2020) and other substrates(Adam et al., 2007; Rotem and Schuldiner, 2004; Soskine et al., 2004). The electrophysiology traces are very similar for WT EmrE and EmrE_3_ (Figure 1B). Measurements of peak capacitive currents as a function of substrate concentration were fit to the Michaelis-Menten equation, yielding K_m_ values for EmrE_3_ within 2-fold of those measured for WT EmrE (Figure 1B, Figure 1 – Figure Supplement 2). Microscale thermophoresis experiments show that EmrE_3_ binds monobody L10 with a K_d_ of 850 nM (Figure 1C, Figure 1 – Figure Supplement 3), indicating that these small modifications at surface exposed residues were sufficient to create a monobody binding site. Similar to our observation for Gdx-Clo(Kermani et al., 2020), addition of saturating L10 monobody (10 μM) depresses transport currents mediated by EmrE_3_ by about 40% but does not altogether inhibit substrate transport (Figure 1D, E). Currents are fully restored upon subsequent incubation with monobody-free solution. Thus, EmrE_3_ is functionally equivalent to WT EmrE, is capable of binding monobody L10, and retains function when this monobody is bound.

### Structure of EmrE_3_ without ligand at pH 5.2

When combined with monobody L10, EmrE_3_ crystallized and diffracted to a maximum resolution of 2.9 Å. The crystallization conditions differed from those used for the Gdx-Clo/monobody complex, but the space group, C121, and approximate dimensions of the unit cell were the same(Kermani et al., 2020). We solved the structure using molecular replacement, with the L10 monobodies and the first three helices of each Gdx-Clo monomer as search models. After phasing, loop 3 and helix 4 were built into the experimental density followed by iterative rounds of refinement (Figure 2A, Table 1, Figure 2-Figure Supplement 1A, B). The model was validated by preparing a composite omit map in which 5% of the atoms in the model were removed at a time (Terwilliger et al., 2008)(Figure 2 — Figure Supplement 1C, D). Our EmrE_3_ model corresponds well with the composite omit maps, suggesting that model bias introduced by using Gdx-Clo as a molecular replacement search model does not unduly influence our model of EmrE_3_.

**Figure 2.**
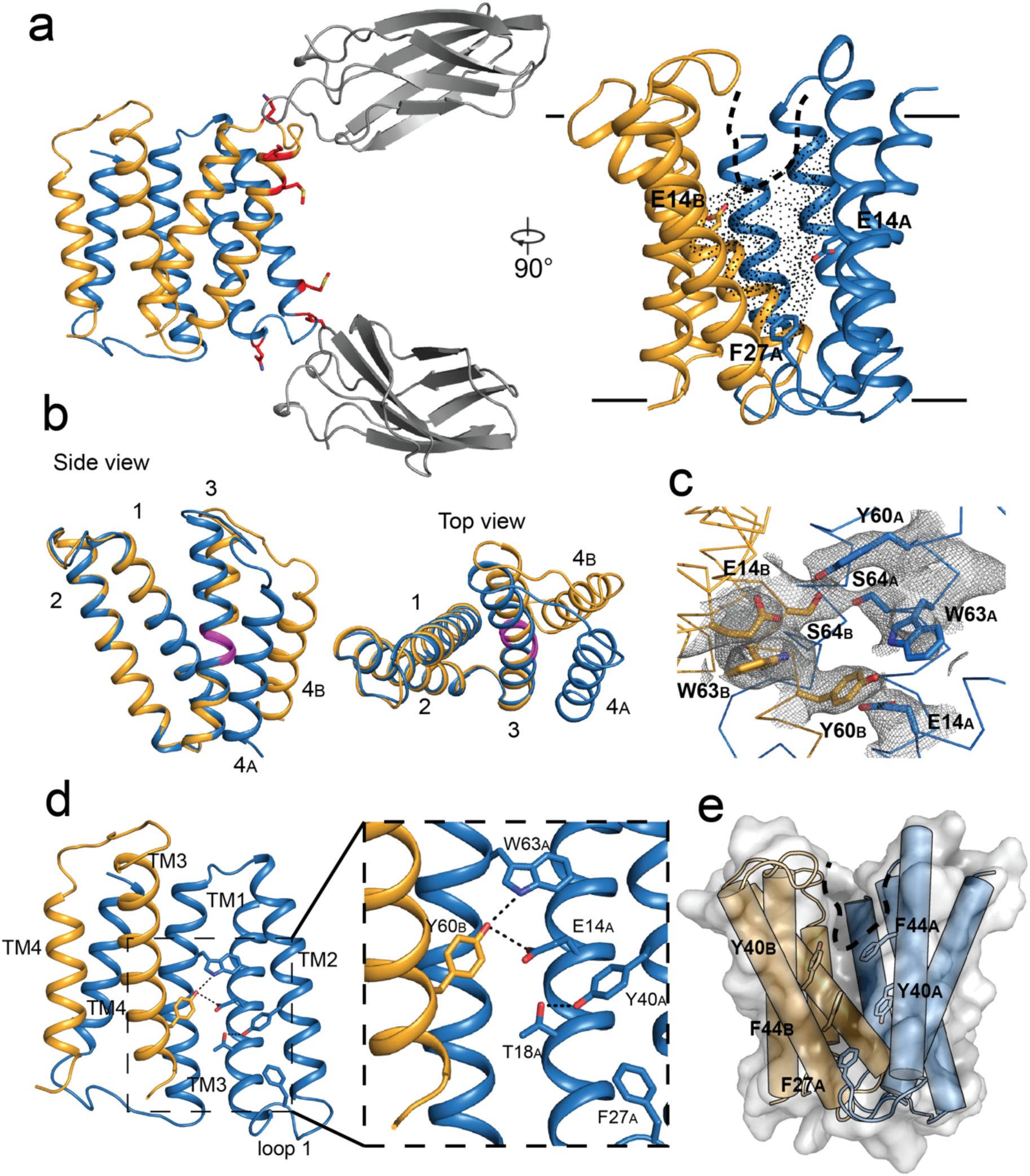
Crystal structure of EmrE_3_. A. Subunits A and B are shown in blue and orange, respectively, and monobody L10 is shown in gray. In the left panel, mutated residues E25N, W31I, V34M are shown in red with sidechain sticks. In the right panel, the monobodies are removed for clarity. E14_A_, E14_B_, and F27_A_ are shown as sticks, and the aqueous accessible region of the transporter is indicated with dots. Approximate membrane boundaries are shown as solid lines, and the boundary of the membrane portal is shown as a dashed line. B. A (blue) and B (orange) subunits of EmrE_3_, aligned over residues 1-63. The GVG fulcrum sequence in TM3 is colored in magenta. C. S64 and surrounding sidechains with 2mF_o_-DF_c_ density shown as gray mesh (contoured at 1.0 *σ* within 2 Å of selected residues). D. Y60_B_ hydrogen bonding network. EmrE dimers are shown with TM1 and TM2 of subunit B (orange) removed for clarity. Lower panels show zoomed in view. In each view, interactions within hydrogen bonding distance and geometry are shown as dashed lines. E. Surface rendering of EmrE_3_. TM2 sidechains that line the portal are shown as sticks.

**Table 1.** Data collection, phasing and refinement statistics for EmrE and Gdx-Clo complexes

|  | EmrE<sub>3</sub>/L10/MeTPP<sup>+</sup> | EmrE<sub>3</sub>/L10/TPP<sup>+</sup> | EmrE<sub>3</sub>/L10/harmane |
| --- | --- | --- | --- |
| **Data collection** |  |  |  |
| Space group | C121 | C121 | C121 |
| Cell dimensions |  |  |  |
| *a*, *b*, *c* (Å) | 141.17, 50.87, 110.79 | 140.71, 50.14, 110.28 | 145.7, 51.83, 114.95 |
| $\alpha$, $\beta$, $\gamma$ (°) | 90, 92.69, 90 | 90, 93.45, 90 | 90, 92.67, 90 |
| Resolution (Å) | 70.5-3.22 (3.42-3.22) | 70.2-3.36 (3.62-3.36) | 114.8-3.75 (4.37-3.75) |
| Ellipsoidal Resolution Limit (best/worst)<sup>a</sup> | 3.22/4.33 | 3.36/5.1 | 3.75/6.34 |
| % Spherical Data Completeness<sup>a</sup> | 69.0 (20.9) | 54.5 (13.6) | 44.0 (10.1) |
| % Ellipsoidal Data Completeness<sup>a</sup> | 88.6 (80.1) | 84.1 (78.5) | 82.7 (65.7) |
| *R*<sub>merge</sub><sup>a</sup> | 0.152 (0.656) | 0.349 (1.053) | 0.365 (0.752) |
| *R*<sub>meas</sub><sup>a</sup> | 0.166 (0.707) | 0.384 (1.15) | 0.396 (0.817) |
| CC<sub>1/2</sub> | 0.967 (0.861) | 0.779 (0.610) | 0.992 (0.862) |
| Mn *I* / $\sigma I$<sup>a</sup> | 10.4 (2.7) | 4.0 (1.8) | 7.4 (2.3) |
| Multiplicity<sup>a</sup> | 6.6 (7.1) | 5.9 (6.2) | 6.7 (6.6) |
| **Refinement** |  |  |  |
| Resolution (Å) | 55.3 - 3.22 | 55.0-3.36 | 60.2-3.91 |
| No. reflections | 8025 | 6097 | 3347 |
| *R*<sub>work</sub> / *R*<sub>free</sub> | 29.4 / 33.4 | 29.0/31.4 | 34.2/34.4 |
| Ramachandran Favored | 89.4 | 89.6 | 90.9 |
| Ramachandran Outliers | 1.9 | 1.9 | 2.4 |
| Clashscore | 11.8 | 13.6 | 8.4 |
| R.m.s. deviations |  |  |  |
| Bond lengths (Å) | 0.003 | 0.003 | 0.003 |
| Bond angles (°) | 0.70 | 0.68 | 0.60 |
| Coordinates in Protein Databank | 7SSU | 7SV9 | 7SVX |

**Table 1. Data collection, phasing and refinement statistics for EmrE and Gdx-Clo complexes (continued from previous page)**
|  | EmrE<sub>3</sub>/L10/methyl<br>viologen | EmrE<sub>3</sub>/L10, pH 5.2 | Gdx-Clo/L10, pH 5.0 | EmrE<sub>3</sub>/L10/BM<sub>3</sub>A<sup>+</sup> |
| --- | --- | --- | --- | --- |
| **Data collection** |  |  |  |  |
| Space group | P1 | C121 | P1 | C121 |
| Cell dimensions |  |  |  |  |
| *a*, *b*, *c* (Å) | 50.91, 75.07, 111.43 | 140.64, 49.85, 109.83 | 49.70, 74.32, 107.43 | 140.18, 50.12, 110.73 |
| $\alpha$, $\beta$, $\gamma$ (°) | 92.03, 90.33, 109.20 | 90, 93.75, 90 | 93.56, 89.71, 109.92 | 90, 92.79, 90 |
| Resolution (Å) | 70.8-3.13 (3.41-3.13) | 70.2-2.85(3.16-2.85) | 107.2-2.32 (2.67-2.32) | 70.50-3.22 (3.42-3.22) |
| Ellipsoidal Resolution Limit (best/worst)<sup>a</sup> | 3.13/4.50 | 2.85/3.72 | 2.32/3.55 | 3.22/4.33 |
| % Spherical Data Completeness<sup>a</sup> | 52.0 (11.1) | 62.0 (12.0) | 41.9 (6.0) | 69.0 (20.9) |
| % Ellipsoidal Data Completeness<sup>a</sup> | 82.0 (72.3) | 87 (62.6) | 80.3 (45.6) | 88.6 (80.1) |
| *R*<sub>merge</sub><sup>a</sup> | 0.123 (0.697) | 0.118 (1.85) | .089 (0.4) | 0.152 (0.656) |
| *R*<sub>meas</sub><sup>a</sup> | 0.144 (0.814) | 0.129 (1.99) | .104 (0.465) | 0.166 (0.707) |
| CC<sub>1/2</sub> | 0.939 (0.629) | 0.994 (0.366) |  | 0.967 (0.861) |
| Mn *I* / σ<sup>a</sup> | 7.7 (1.5) | 9.5 (1.2) | 6.5 (2.8) | 10.4 (2.7) |
| Multiplicity<sup>a</sup> | 3.7 (3.8) | 6.4 (7.1) | 3.8 (3.8) | 6.6 (7.1) |
| **Refinement** |  |  |  |  |
| Resolution (Å) | 32.9-3.13 | 35.2-2.85 | 35.5-2.32 | 55.3-3.91 |
| No. reflections | 14,194 | 11,149 | 26,026 | 5040 |
| *R*<sub>work</sub> / *R*<sub>free</sub> | 30.0/33.1 | 30.7/32.7 | 25.1/29.5 | 33.0/36.7 |
| Ramachandran Favored | 89.1 | 91.0 | 92.9 | 88.7 |
| Ramachandran Outliers | 2.6 | 1.9 | 1.5 | 2.7 |
| Clashscore | 16.8 | 8.6 | 10.4 | 14.9 |
| R.m.s. deviations |  |  |  |  |
| Bond lengths (Å) | 0.004 | 0.003 | 0.004 | 0.004 |
| Bond angles (°) | 0.82 | .65 | 0.70 | 0.67 |
| Coordinates in Protein Databank | 7MGX | 7MH6 | 7SZT | 7T00 |

The structure of the EmrE_3_/L10 complex (Figure 2A) shows an antiparallel EmrE_3_ dimer bound to two monobodies in slightly different orientations via the loop 1 residues. The crystal packing is similar to Gdx-Clo, with the majority of contacts mediated by monobody. The introduced E25N sidechain of EmrE_3_ is within hydrogen bonding distance of a tyrosine sidechain contributed by the monobody, and W31I contributes to a hydrophobic patch of the transporter/monobody interface. These interactions are homologous to those observed for the Gdx-Clo/L10 complex. The third mutant sidechain of EmrE_3_, V34M, does not interact with monobody in this structure, and therefore might not be necessary for monobody binding to EmrE_3_.

In our EmrE_3_ model, the positions of the helices are consistent with existing electron microscopy maps of EmrE (∼8 Å resolution) (Ubarretxena-Belandia et al., 2003) (Figure 2 – Figure Supplement 2A). Compared with a previous MD model based on that EM data(Vermaas et al., 2018), our current EmrE_3_ crystal structure has a C*_α_*RMSD of 2.5 Å, with close correspondence of residues that contribute to the substrate binding pocket (Figure 2 – Figure Supplement 2B). Although EmrE_3_ has high structural similarity to Gdx-Clo (C*_α_*RMSD 1.2 Å for the dimer), the structures display clear differences in subunit packing. Relative to Gdx-Clo, in EmrE_3_ helices 1-3 of the A subunit, which line the binding pocket, are each displaced by 1.5 – 2.5 Å (Figure 2 – Figure Supplement 2C). These shifts slightly expand the aqueous cavity of EmrE_3_ relative to Gdx-Clo.

As in Gdx-Clo, the two monomers adopt different structures. Monomers A and B differ from each other in the relative orientation of their two domains about a fulcrum at the conserved GVG motif in helix 3 (residues 65-67; Figure 2B). The domains are comprised of the first 65 residues (helices 1, 2, and helix 3 until the GVG motif) and the last 40 residues (the C-terminal half of helix 3 and helix 4). The observed domain architecture is in accord with the proposed conformational swap of two structurally distinct monomers(Morrison et al., 2012). The residue S64 is positioned immediately before the GVG fulcrum, at the boundary of domain 1 and domain 2 for each EmrE_3_ subunit. In the crystal structure, the S64 sidechains contributed by the two subunits are within hydrogen bonding distance and geometry, with strong contiguous electron density between them (Figure 2C). Due to the antiparallel architecture, the outward- and inward-facing conformations of the transporter are expected to be structurally identical and related by 2- fold symmetry about an axis parallel to the plane of the membrane(Morrison et al., 2012). Thus, the S64 interaction should be preserved when the transporter is open to the opposite side of the membrane; we therefore imagine that the S64 sidechains remain hydrogen bonded to each other during the entire transport cycle, forming the pivot point around which the conformational change occurs.

In the absence of ligand, EmrE_3_ possesses a deep, spacious aqueous pocket that is accessible from one side of the membrane (Figure 2A). The E14 sidechains contributed by both subunits define the edges of this binding pocket. E14 is invariant in the SMR family and essential for binding both substrate and protons(Yerushalmi and Schuldiner, 2000). The present crystals formed at pH 5.2, at which both E14 sidechains are expected to be protonated (Li et al., 2021; Morrison et al., 2015). There is a small, spherical density in the vestibule between W63_B_ and E14_A_ that is consistent with a water molecule, although no other ordered water molecules are visible at this resolution (Figure 2 – Figure Supplement 3). The cross-subunit interaction between Y60_B_ and E14_A_ proposed by Vermaas *et al*. is observed (Figure 2D). A conserved hydrogen bond acceptor, T17_A_, is located one helical turn down from E14_A_ and engaged in an intrasubunit interaction with Y40_A_ (Figure 2D).

As in Gdx-Clo, the TM2 helices splay apart on the open side of the transporter, defining a portal from the membrane to the substrate binding site that is lined with hydrophobic sidechains (Figure 2E). This portal may play a dual role, rearranging to allow alkyl substituents to reside in the membrane during the transport cycle, as well as providing the opportunity for hydrophobic drugs to diffuse laterally from the membrane into the substrate binding site. Aromatic residues contributed by loop 1_A_, including the highly conserved F27 sidechain, are wedged between the hydrophobic sidechains lining helices 2_A_ and 2_B_, sealing the closed side of the transporter (Figure 2E).

### Structures of substrate-bound EmrE_3_

To understand how different substrates interact with EmrE, we screened a variety of transported compounds in crystallization trials at pH values ≥6.5, where the E14 sidechains are expected to be deprotonated, favoring binding of the positively charged substrates. We were able to obtain diffracting crystals in the presence of five structurally diverse compounds transported by EmrE: monovalent planar aromatic harmane (3.8 Å), divalent planar aromatic methyl viologen (3.1 Å), quaternary phosphoniums tetraphenylphosphonium (TPP^+^; 3.4 Å) and methyltriphenylphosphonium (MeTPP^+^; 3.2 Å), and quaternary ammonium benzyltrimethylammonium (3.9 Å) (Table 1). We were unable to generate crystals that diffracted to high resolution in the presence of metformin, benzalkonium, cetyltrimethylammonium, or octylguanidinium. Phases were determined using molecular replacement with the pH 5.2 EmrE_3_ monomers and L10 monobodies as search models. Although the crystallization conditions differed for each substrate, the TPP^+^-, MeTPP^+^-, benzyltrimethylammonium-, and harmane-bound proteins crystallized in the same unit cell as proton-bound EmrE_3_, with one copy of the EmrE_3_/L10 complex in the asymmetric unit. The methyl viologen-bound protein crystallized in P1 with two pseudosymmetric copies of the EmrE3/L10 complex in the asymmetric unit, organized in the same relative orientation as individual complexes in the C121 crystal form.

Since Gdx-Clo and EmrE_3_ were both accommodated in this crystal lattice despite differences in the tilt and packing of helices 1, 2, and 3, we expect that small 1-2 Å substrate-dependent movements in the backbone of EmrE_3_ would also be tolerated within this crystal lattice. However, in all five substrate-bound structures, the transmembrane helices and loops 1 and 2 conform almost perfectly to the pH 5.2 structure (C*_α_*RMSD = 0.5-0.65 Å), suggesting that the observed backbone conformation is the lowest energy state for both the substrate- and proton-bound transporter. Loop 3 is poorly ordered and adopts a different conformation in each of the structures in which it is resolved well enough to model.

For all substrate-bound structures, the maps show positive densities between the substrate-binding E14 residues, including a four-lobed density for TPP^+^, a three-lobed density for MeTPP^+^, and oblong densities for the harmane and the methyl viologen structures. We modelled the corresponding substrates into each of these densities (Figure 3A, B). All five drugs are bound at the bottom of the aqueous cavity, in overlapping positions at the midpoint of the membrane. In the two copies of the methyl viologen-bound transporter, the drug is bound in different (but overlapping) positions (Figure 3A, Figure 3 – Figure Supplement 1). For all substrates, the center of mass is poised midway between the E14 residues. To different extents, the substrates also interact with the protein’s aromatic residues via ring stacking, especially Y60 and W63.

**Figure 3.**
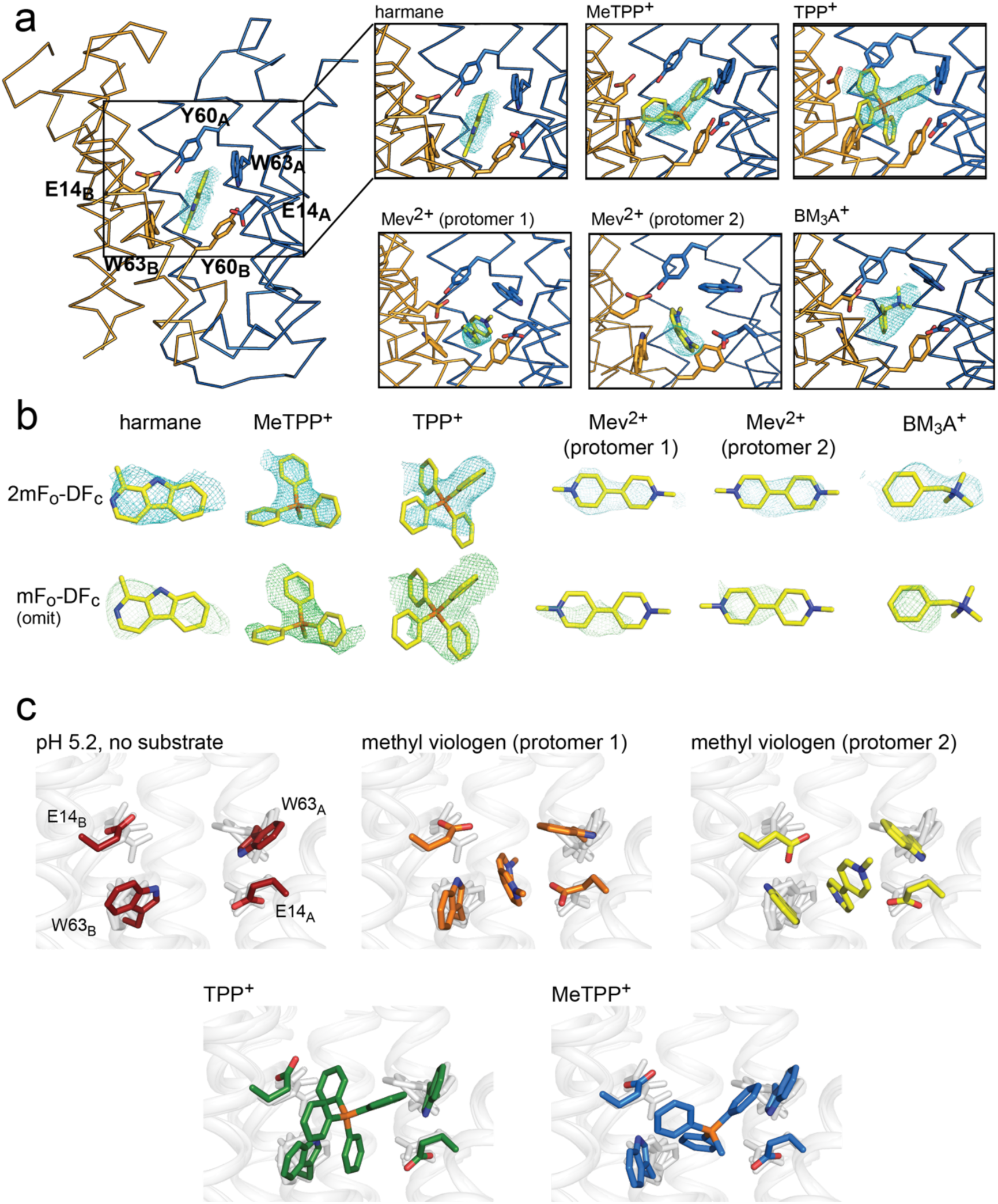
Substrate binding to EmrE_3_. A. Structures are shown in ribbon representation, with sidechains E14, W63, and Y60 shown as sticks. All panels are zoomed and oriented the same. 2mF_o_-DF_c_ maps (carved 2 Å around each substrate) are shown as cyan mesh. Maps are contoured at 1*σ* for harmane and 1.2*σ* for MeTPP^+^, TPP^+^, methylviologen, and benzyltrimethylammonium (BM_3_A^+^). B. Top row: Substrate structures and 2mF_o_-DF_c_ maps from the panels in A, individually rotated to view each substrate. Bottom row: mF_o_-DF_c_ substrate omit maps shown as green mesh. Omit maps are contoured at 1.8*σ* for harmane and 2*σ* for MeTPP^+^, TPP^+^, methylviologen, and BM_3_A^+^. C. Comparison of E14 and W63 positions in each substrate-bound structure. Individual panels show substrate, E14, and W63 from indicated structure in color aligned with the other four structures, which are rendered in light gray.

Comparison of these structures permitted evaluation of the specific orientations of the sidechains that line the substrate binding site (Figure 3C). The harmane- and benzyltrimethylammonium-bound structure was excluded from this analysis because, at 3.8 – 3.9 Å resolution, we were not as confident about interpreting subtle changes in sidechain orientation. For the other substrates (methyl viologen, TPP^+^, and MeTPP^+^), this comparison showed that binding site sidechains, especially E14 and W63, adopt different rotamers, thus accommodating the differently sized substrates. For example, the carboxylate of E14_B_ is displaced by 2.5 Å when the bulky quaternary phosphonium TPP^+^ is bound, compared to its position when the planar methyl viologen occupies the binding site. Likewise, the position of the W63_A_ indole ring rotates over approximately 80° depending on the substrate that occupies the binding site. To validate these observations, we performed refinements with models in which the position of the W63_A_ or E14_B_ sidechain was adjusted to match its position in the presence of a dissimilar substrate; the resulting difference density demonstrates that these substrate-dependent changes in sidechain rotamer are not due to model bias during the refinement (Figure 3 – Figure Supplement 2, Figure 3 – Figure Supplement 3). Thus, these structures provide a first suggestion of how rotameric movements of EmrE’s charged and aromatic sidechains can change the dimensions of the binding pocket and interact favorably with diverse substrates.

### Structure of Gdx-Clo at pH 5 and comparison to the substrate binding site of EmrE

The overall fold and many of the binding site sidechains are shared between EmrE and Gdx-Clo, yet the two proteins have markedly different substrate selectivity profiles. We therefore sought to analyze how molecular interactions among binding site residues might explain the different substrate selectivity for EmrE and Gdx-Clo. Previous structures of Gdx-Clo were solved at pH ≥7.5 in complex with substituted guanidinyl compounds(Kermani et al., 2020). In order to compare the substrate binding sites of Gdx-Clo and EmrE_3_ in equivalent states, we solved a new structure of Gdx-Clo at pH 5.0, which is close to the value for the present low pH EmrE_3_ structure, pH 5.2 (Table 1, Figure 4 – Figure Supplement 1A). Both transporters are likely proton-bound at this pH, minimizing differences in sidechain positioning that might stem from interactions with bound substrate. This new structure of proton-bound Gdx-Clo, which is resolved to 2.3 Å, is highly similar to the structure of substrate-bound Gdx-Clo (PDB: 6WK8), with only a local change in the rotamer of the central glutamate E13_B, Clo_ (Figure 4 – Figure Supplement 1B).

A comparison of the low-pH EmrE_3_ and Gdx-Clo structures reveals conspicuous differences in the hydrogen bond network within the binding cavity (Figure 4A, B), despite the conservation of many key residues. In Gdx-Clo, S42_Clo_ participates in the stack of alternating hydrogen bond donors and acceptors (W16_Clo_/E13_Clo_/S42_Clo_/W62_Clo_) that fixes the position of the central Glu, E13_Clo_. Although the analogous serine (S43_EmrE_) is present in EmrE, it is not playing an analogous role. A 1.5 Å displacement in helix 2 has distanced this Ser from the other sidechains in the binding pocket, beyond hydrogen bonding distance with W63_EmrE_. Instead, S43_EmrE_ is rotated away from the aqueous cavity and the central E14_EmrE_ residues. Despite strict conservation of this serine among the Gdx subtype, mutation to alanine is common among the Qacs (Figure 4C). In lieu of an interaction with S43_EmrE_, both W63_EmrE_ sidechains in EmrE adopt different rotamers compared to their counterparts in Gdx-Clo. W63_A, EmrE_ is oriented so that its indole NH is within H-bonding distance of Y60_B, EmrE_, although the angle between the H-bond donor and acceptor is suboptimal, approximately 30° off normal.

**Figure 4.**
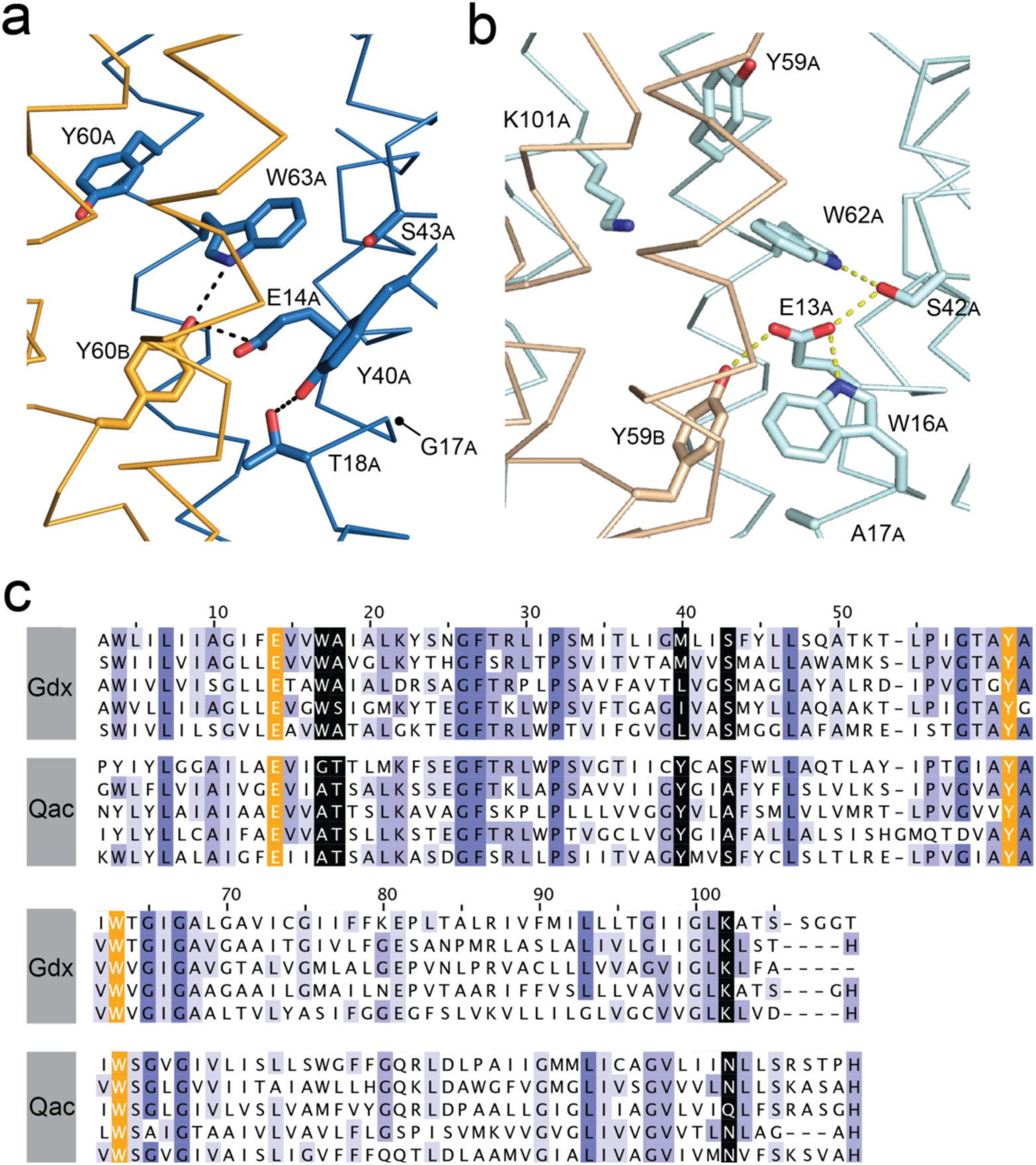
Structure and sequence alignment of substrate binding site residues in Qac and Gdx subtypes. A. Substrate binding site in EmrE_3_, with subunit B in orange and subunit A in blue. B. Substrate binding site in Gdx-Clo, with subunit B in wheat and subunit A in pale cyan (PDB: 6WK8). For panels A and B, the proteins are shown in the same orientation. Note that residue numbering is offset by one in Gdx-Clo. Potential hydrogen bonds are shown as dashed lines. C. Sequence alignments of five representative Gdx proteins (from top to bottom: *Clostridiales* bacterium oral taxon 876, *E. coli*, *Micromonospora*, *Streptomyces tsukubensis*, and *Leifsonia aquatica*) and five representative Qac proteins (from top to bottom: *E. coli*, *Klebsiella pneumoniae*, *Pseudomonas aeruginosa*, *Mycobacterium bovis*, and *Bordetella avium*). Sequence numbering corresponds to EmrE. Sequences are colored according to sequence conservation (shades of blue). Residues that contribute to the binding pocket and that are conserved in both the Qac and Gdx subtypes are highlighted in orange. Residues that contribute to the binding pocket and that differ between the Qac and Gdx subtypes are highlighted in black.

The fourth residue from Gdx-Clo’s H-bond stack, W16_Clo_, is universally conserved in Gdx proteins, but replaced with a glycine or alanine in the Qacs (G17 in EmrE). There is no equivalent H-bond donor to the central Glu in EmrE. Instead, the sidechain Y40_EmrE_ occupies this space, but interacts with T18_EmrE_ located one helical turn away from E14_EmrE_. This pair, Y40_EmrE_ and T18_EmrE_, is highly conserved among the Qacs, and variable and typically hydrophobic in Gdx proteins. In Gdx-Clo, the corresponding positions are M39_Clo_ and A17_Clo_. This trio of correlated positions (W16_Clo_/G17_EmrE_, A17_Clo_/T18_EmrE_, and M39_Clo_/Y40_EmrE_) in the substrate binding site are among the main features that differentiate the Gdx and Qac subtypes in sequence alignments (Figure 4C).

Y60_A, EmrE_ also adopts a different orientation in EmrE relative to the position of the analogous Tyr, Y59_Clo_ in Gdx-Clo. Rather than extending out of the binding pocket towards the exterior solution, as it does in Gdx-Clo, Y60_A, EmrE_ is pointed down towards the S64_EmrE_ diad. This rotamer would not be possible in Gdx-Clo, since this space is occupied by K101_Clo_ instead, which extends from the C-terminal end of helix 4 and points down into the substrate binding pocket towards the glutamates. K101_Clo_ is completely conserved in the Gdx subtype.

The overall picture that emerges from this comparison of the Gdx-Clo and EmrE structures is that the two proteins share many binding site residues but differ in the relative organization of these residues. In Gdx-Clo, E13_Clo_, S42_Clo_, Y59_Clo_, and W62_Clo_ are constrained in a highly organized H-bond network. In EmrE, residues peripheral to the binding site have encroached on these positions, disrupting the network and reducing the number of protein hydrogen bond partners for each of these conserved sidechains.

### EmrE is tolerant of mutations that eliminate hydrogen bonding in the binding pocket

Based on structural comparison of the Gdx-Clo and EmrE binding pockets, we hypothesize that even for conserved residues in the binding pocket, the importance of hydrogen bonding is diminished in EmrE relative to Gdx-Clo. To probe this, we performed a head-to-head comparison of SSM currents mediated by EmrE and Gdx-Clo proteins with mutations at three conserved positions adjacent to the functionally essential central Glu: Y59F_Clo_/Y60F_EmrE_, S42A_Clo_/S43A_EmrE_, and W62F_Clo_/W63F_EmrE_ (Figure 5, Table 2). All six mutant transporters were expressed at near-WT levels and monodisperse by size exclusion chromatography. For EmrE mutants, we tested transport of 2 mM PheGdm^+^ or 2 mM TPA^+^, and for Gdx-Clo, we tested transport of its native substrate, 1 mM Gdm^+^. For all experiments, substrate concentration was ∼4-fold higher than the transport K_m_.

**Figure 5.**
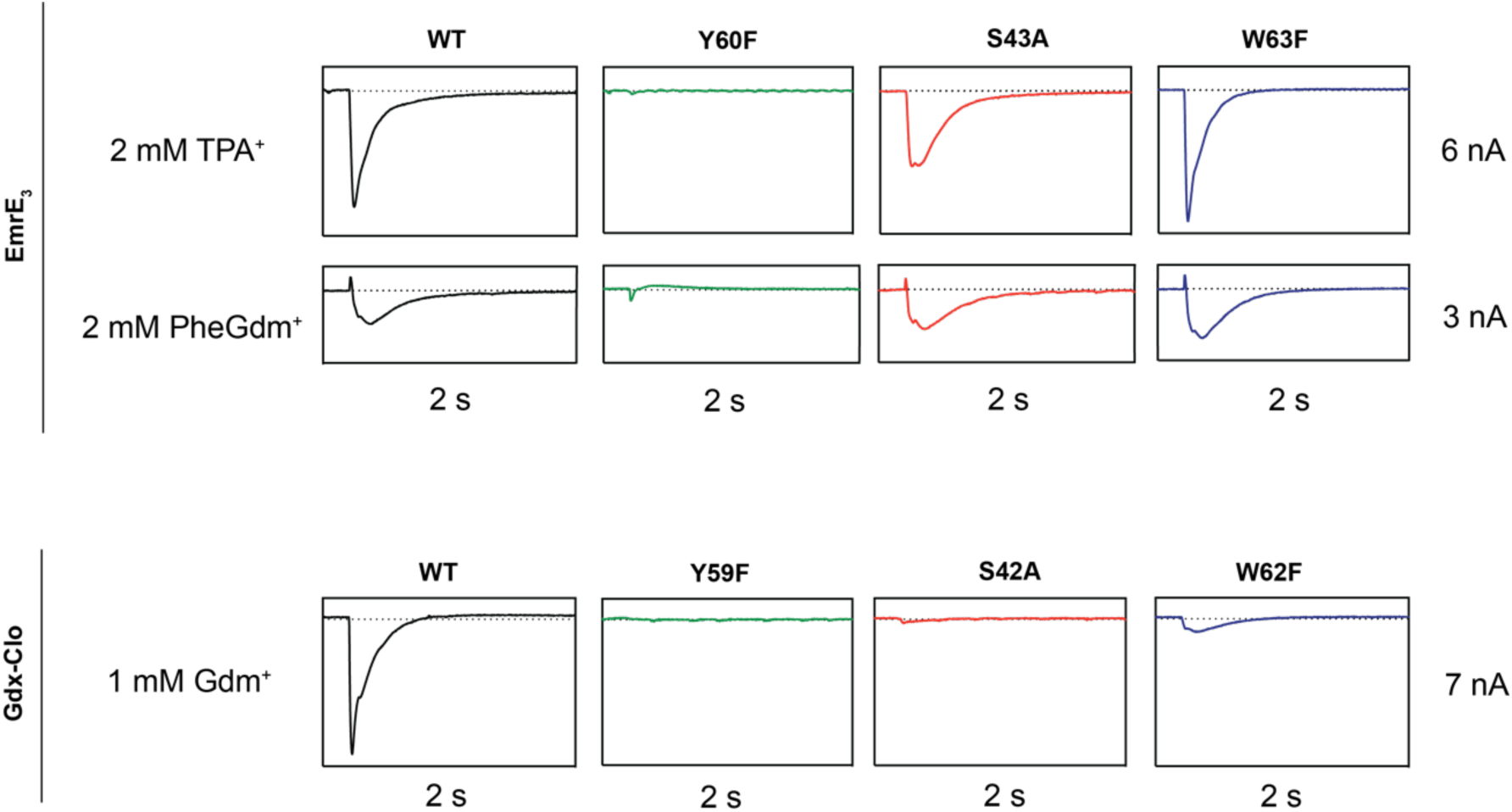
Representative SSM electrophysiology recordings for EmrE_3_ and Gdx-Clo mutants. For EmrE_3_, PheGdm^+^ and TPA^+^ traces are from the same sensor and shown on the same scale. Vertical box edges are 3 nA for PheGdm^+^ traces, and 6 nA for TPA^+^ traces. For Gdx-Clo, vertical box edges are 7 nA. Horizontal box edges are 2 s for all traces. Dashed line represents the zero-current level. Traces are representative of currents from three independently prepared sensors and two independent biochemical preparations. Peak current values for all replicates are reported in Table 2. Note that because there is some sensor-to-sensor variation in liposome fusion, comparisons of current amplitude among the mutants are qualitative.

**Table 2.** SSM electrophysiology peak currents (nA) for EmrE_3_ and Gdx-Clo mutants summarized by experimental replicate.

|  | Prep 1/Sensor 1 |  | Prep 1/Sensor 2 |  | Prep 2/Sensor 1 |  |
| --- | --- | --- | --- | --- | --- | --- |
|  | TPA<sup>+</sup> | pheGdm<sup>+</sup> | TPA<sup>+</sup> | pheGdm<sup>+</sup> | TPA<sup>+</sup> | pheGdm<sup>+</sup> |
| No protein | 0 | 0 | 0 | 0 | 0 | 0 |
| WT | -4.8 | -1.4 | -4.1 | -1.2 | -3.9 | -1.0 |
| Y60F | -0.14 | -0.03 | -0.05 | -0.07 | -0.04 | -0.05 |
| S43A | -3.7 | -1.6 | -3.9 | -1.3 | -3.2 | -1.2 |
| W63F | -5.4 | -2.0 | -4.6 | -1.5 | -4.0 | -1.0 |

**Gdx-Clo**
|  | Prep 1/Sensor 1 |  | Prep 1/Sensor 2 | Prep 2/Sensor 1 |
| --- | --- | --- | --- | --- |
|  | Gdm<sup>+</sup> |  | Gdm<sup>+</sup> | Gdm<sup>+</sup> |
| No protein | 0 |  | 0 | 0 |
| WT | -6.3 |  | -6.7 | -6.3 |
| Y60F | 0.04 |  | 0.007 | 0.6 |
| S43A | -0.15 |  | -0.15 | -0.15 |
| W63F | -0.60 |  | -0.33 | -0.30 |

In line with its proposed role as a conformational switch(Kermani et al., 2020; Vermaas et al., 2018), no currents were observed when the binding site Tyr (Y59_Clo_/Y60_EmrE_) was mutated in either protein. This result recapitulates results from prior radioactive uptake studies of both mutants(Kermani et al., 2020; Rotem et al., 2006). It also establishes a dead-transporter control for our SSM electrophysiology assays. We likewise find that Gdx-Clo does not tolerate perturbation to its hydrogen bond stack. Although neither S42A_Clo_ nor W62F_Clo_ directly bind Gdm^+^, both mutations eliminate Gdm^+^ currents in SSM electrophysiology assays. In contrast, EmrE_3_ was relatively indifferent to the S43A_EmrE_ and W63F_EmrE_ mutations, with robust currents evoked by both TPA^+^ and PheGdm^+^.

This result for S43_EmrE_ reinforces the structural suggestion that the serine’s functional role in the Gdx transporters is not conserved in the Qac subtype, and is also in agreement with prior transport and resistance assays that showed that S43_EmrE_ modulates substrate specificity in EmrE, but is not required for transport function (Brill et al., 2015; Wu et al., 2019). The observation of robust transport by W63F_EmrE_ is more surprising, since numerous biochemical experiments have demonstrated that W63_EmrE_ mutants do not bind or transport aromatic substrates(Amadi et al., 2010; Elbaz et al., 2005; Wu et al., 2019). To our knowledge, the consequences of W63_EmrE_ mutation have not been previously investigated for non-aromatic substrates in biochemical assays. Our SSM electrophysiology results suggest that maintaining a hydrogen bond donor at W63_EmrE_ is not essential, and that the conservation of W63_EmrE_ is not a mechanistic requirement for EmrE transport, but is instead a determinant of aromatic substrate specificity. In agreement with this interpretation, bacterial growth assays have shown that W63_EmrE_ mutants retain resistance to non-aromatic biocides(Saleh et al., 2018).

## Discussion

In this work, we describe substrate-and proton-bound crystal structures of the *E. coli* SMR transporter EmrE, which is wildtype except for three conservative, functionally neutral mutations that enable monobody binding, and thus, crystallization. Functional assays show that the engineered protein, EmrE_3_ behaves like wildtype, and that the transporter remains capable of transport in the presence of monobody. Below, we discuss the crystallization strategy, we evaluate differences between our crystal structures and a recent NMR-derived model of EmrE(Shcherbakov et al., 2021), and discuss the implications of our structures for understanding substrate polyspecificity by EmrE.

### The application of multipurpose chaperones for crystallization

The minimal monobody binding interface permitted a crystallization chaperone developed for Gdx-Clo to be repurposed for binding and crystallization of a new target with structural homology, but only 35% sequence identity to the original, streamlining the structural characterization process. Given the similarity of this loop among diverse SMR proteins, we think that this approach would likely facilitate the structural characterization of any target within the SMR family. Such general adapters and chaperones to facilitate structural biology have been described before for various targets (Dutka et al., 2019; Koldobskaya et al., 2011; McIlwain et al., 2021; Mukherjee et al., 2020). Although identification of a general SMR monobody was not the original intent of the monobody selection, in cases where multiple homologous targets have been identified, variants with identical or near-identical epitopes could be generated, and binders with broad utility could presumably be selected for. Especially in the case of bacterial proteins, in which there are many clinically relevant homologues from many diverse species, such general structural biology approaches hold particular promise to facilitate molecular characterization of membrane protein targets.

The monobody chaperones mediate most of the crystal contacts, permitting Gdx-Clo and EmrE to crystallize in a nearly identical unit cell, despite some structural differences, including 1-2 Å displacements of helices that contribute to the binding pocket. Although it is a misconception that crystallization chaperones can “force” the transporter into a non-native, high-energy conformation (Koide, 2009), it is plausible that the monobody chaperones recognize a less-prevalent conformation, and kinetically trap the transporter in a minority state within the native conformational ensemble. Because these monobodies were not selected against EmrE, but against a different homologue from the SMR family, this is a possibility that should be considered. However, two lines of evidence disfavor the possibility that the monobody-bound state is aberrant. First, we showed that monobody binding has only a minor effect on transport function, and second, our model corresponds closely to the helix density in the EM dataset, which was obtained without exogenous binding proteins(Ubarretxena-Belandia et al., 2003).

### Comparison to the NMR model of EmrE S64V

An NMR-based model of the “slow-exchanging” EmrE mutant S64V was recently published(Shcherbakov et al., 2021). S64V binds substrate with similar affinity as wildtype, but the rate of conformational exchange is about an order of magnitude slower(Wu et al., 2019). This model was computed based on chemical shift measurements and distance restraints between the protein backbone and a fluorinated substrate tetrafluorophenylphosphonium (F-TPP^+^). Although our present crystal structures agree with the NMR model in general aspects, such as the antiparallel topology, there are also notable differences in the global conformation, with an overall RMSD of 2.3 Å for the two models. Relative to other models of EmrE, including the computational models(Ovchinnikov et al., 2018; Vermaas et al., 2018), the EM *α*-helix model(Ubarretxena-Belandia et al., 2003), and the present crystal structures, in the NMR model the first domain of the A subunit is shifted down in a direction perpendicular to the membrane with respect to the B subunit (Figure 6A). (Note that chain A of the NMR structure is more structurally homologous to chain B of the crystal structure and vice versa. Our designation of chains A and B in the present crystal structure correspond to the A and B chains in previous literature, including SMR family homologue Gdx-Clo(Kermani et al., 2020), the low resolution structures of EmrE(Chen et al., 2007; Fleishman et al., 2006), and theoretical EmrE models(Ovchinnikov et al., 2018; Vermaas et al., 2018).) This difference in subunit packing is accompanied by subtle differences in the tilts of the helices (Figure 6B). In the NMR structure, helix 2_A_ and 2_B_ become more parallel, and the gap between them is narrowed, reducing membrane access to the binding site via the portal.

**Figure 6.**
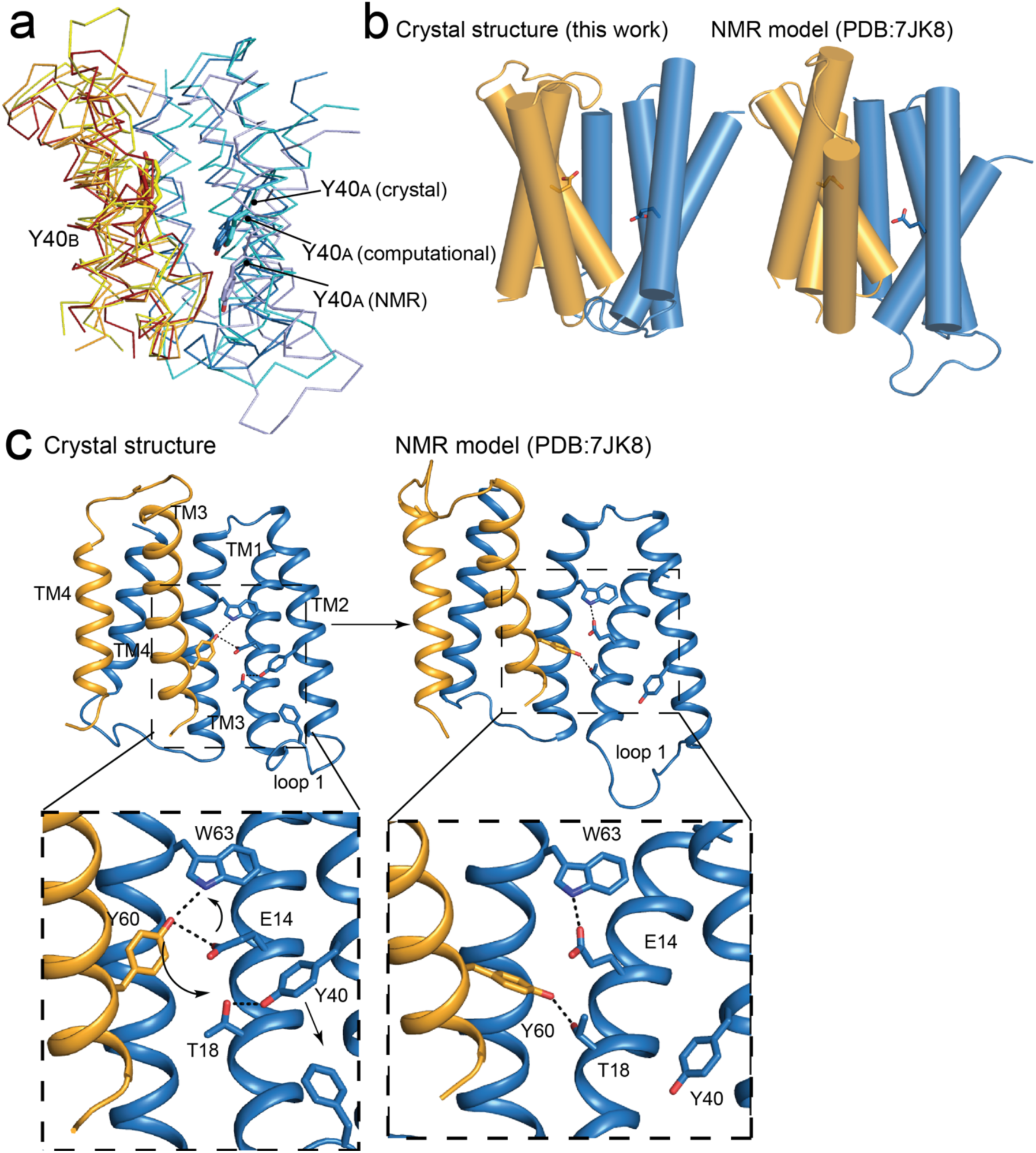
Comparisons of NMR and crystallography models of EmrE. A. Overlay of crystallography (orange/blue), computational (yellow/cyan; (Vermaas et al., 2018)) and NMR (dark red/pale blue; (Shcherbakov et al., 2021)) models, aligned over the B subunit. Y40 sidechain sticks are show as landmarks. B. Side-by-side comparison of the crystallography and NMR models, with A subunit in blue and B subunit in orange. E14 sidechains shown as landmarks. C. Comparison of Y60_B_ hydrogen bonding network in the crystal structure (left) and NMR structure (right). EmrE dimers are shown with TM 1 and 2 of subunit B (orange) removed for clarity. Lower panels show zoomed in view. In each view, interactions within hydrogen bonding distance and geometry are shown as dashed lines. Arrows are shown to help visualize sidechain rearrangements between the two structures.

The difference in global conformation of the NMR and crystallography models is supported by a reorganization of the hydrogen bonding network in the substrate binding site (Figure 6C). The heart of this change is a rotameric switch by Y60: In the crystal structures, Y60_B_ participates in a pair of cross-subunit interactions, within coordination distance and geometry of E14_A_ and W63_A_ in the opposite subunit. In the NMR model, the same Y60_B_ sidechain is assigned a different rotamer, its hydroxyl moving 6 Å along helix 1, so that it is now coordinating T18_A_, one helical turn away from E14_A_. The interaction with Y60_B_ has displaced Y40_A_ from its interaction with T18_A_. Helix 2_A_ slides in a direction perpendicular to the membrane so that Y40_A_ now encroaches on the position of F27_A_ at the tip of loop 1, which is packed between helices 2_A_ and 2_B_ in the crystal structure. In the NMR ensemble, the displaced loop 1 is flexible and adopts various conformations. The previous EM maps correspond closely to the present crystallography models (Real space correlation coefficient (RSCC) = 0.67), and are less consistent with the NMR model (RSCC = 0.51; Figure 6 – Figure Supplement 1)(Ubarretxena-Belandia et al., 2003).

The differences in conformation between the crystallography/EM maps and the NMR model are unlikely to be due to membrane mimetic (which is shared for the EM and NMR datasets), the presence of monobodies (the EM data was collected without monobodies), or the S64V mutation used for NMR studies (NMR experiments showed little change in backbone configuration for this mutant(Wu et al., 2019)). It is possible that the different temperatures and biochemical conditions of the NMR and crystallography experiments favor different states in a conformational ensemble. Previous EPR measurements may lend support to this possibility (Dastvan et al., 2016). Those experiments showed that at pH 8, with TPP^+^ bound, EmrE adopts a major conformation consistent with our current crystallography model. But when substrate is removed and the pH dropped to 5.5, EmrE’s conformational ensemble becomes more heterogeneous. The loops disengage and become more flexible, and a population emerges in which the two subunits have adopted a more-symmetric conformation. Perhaps the NMR experiments, which were performed at pH 5.8 (albeit with substrate) reflect that second conformation from the ensemble. Nevertheless, it is also worth noting that our crystallography model is not inconsistent with the backbone chemical shifts measured in bicelles, based on structure-trained predictions of chemical shift(Frank et al., 2015; Xie et al., 2020) (Figure 6-Figure Supplement 2). Future studies will be required to assess the relevance of these different states to the transport mechanism of EmrE.

### Sidechain movements accommodate diverse substrates

EmrE has been studied in great breadth and depth. Full mutagenic scans coupled with growth assays(Amadi et al., 2010; Gutman et al., 2003; Mordoch et al., 1999; Wu et al., 2019), functional assays with reconstituted transporter (reviewed in (Schuldiner, 2009)), and EPR and NMR spectroscopy experiments(Amadi et al., 2010; Banigan et al., 2015; Dastvan et al., 2016; Leninger et al., 2019; Thomas et al., 2018) have all revealed detailed information about the positions that contribute to substrate binding and conformational change, even as the structural details were lacking. Our structure corroborates many of the specific predictions regarding sidechains that contribute to the binding pocket, including the importance of W63 for aromatic packing with the substrate(Elbaz et al., 2005) and the cross-subunit engagement of Y60(Vermaas et al., 2018). Positions that are sensitive to mutation, including E14, T18, Y40, and L47 all line the binding pocket in our structures (Mordoch et al., 1999; Rotem et al., 2006; Saleh et al., 2018; Wu et al., 2019). Our structure also confirms other architectural features proposed from spectroscopic studies, including the deflection of loop 2 sidechain F27_A_ towards the substrate bound in the binding pocket and the positioning of the portal-lining Y40 and F44 sidechains as an access point from the membrane to the substrate binding site(Dastvan et al., 2016).

In addition to substantiating prior EmrE experiments, our structures also provide new molecular insights into the binding of structurally diverse substrates by EmrE. In addition to a substrate-free, pH 5.2 structure, we solved structures of EmrE with methyl viologen, harmane, Me-TPP^+^, TPP^+^, and benzyltrimethylammonium. These compounds have considerable structural differences, but are all accommodated in the EmrE binding site with only sidechain rearrangements.

The closely related, but substantially more selective SMR family member, Gdx-Clo, provides a useful point of comparison to understand why EmrE can interact with this chemically diverse range of compounds. In Gdx-Clo, the substrate-binding glutamate sidechains are constrained by a polarized stack of hydrogen bond donors and acceptors that also includes W16_Clo_, S42_Clo_, and W62_Clo_. This hydrogen bonded network would be disrupted by the rotamerization of either E13_Clo_ or W62_Clo_. We show that in Gdx-Clo, mutations to sidechains that contribute to the hydrogen bond stack seriously impair transport activity.

In contrast, in EmrE, the corresponding residues E14_EmrE_ and W63_EmrE_ are not constrained by such a stack of H-bond donors and acceptors. The current structures and SSM electrophysiology experiments both suggest that, in contrast to Gdx-Clo, a rigid H-bond network is not essential for substrate transport by EmrE, which remains functional when hydrogen bond capacity is eliminated at S43_EmrE_ or W63_EmrE_. Without the stricter geometric constraints imposed by a polarized stack of sidechain hydrogen bond partners, both E14_EmrE_ and W63_EmrE_ have more flexibility to adopt different rotamers. Like a pair of calipers, the E14_EmrE_ sidechains can move farther apart to accommodate large substrates such as quaternary ammoniums, or closer together for flat, aromatic substrates or substrates with small headgroups, like harmane and methyl viologen or singly substituted guanidinyl compounds. Similarly, W63_EmrE_ has the space and flexibility to rotamerize, which can expand or narrow the binding pocket or allow W63_EmrE_ to rotate in order to pack against the aromatic groups of bound substrates. These structural observations are in agreement with numerous prior studies that have demonstrated an important role for W63_EmrE_ in transport of polyaromatic substrates (Amadi et al., 2010; Elbaz et al., 2005; Saleh et al., 2018; Wu et al., 2019). We note that although W63_A, EmrE_ does change position to conform to different substrates, we did not always observe optimal pi stacking geometry between the substrate and the protein’s aromatic residues. Instead, substrate positioning appeared to optimize electrostatic interactions first, with all substrates situated directly between E14_A, EmrE_ and E14_B, EmrE_.

Likewise, many EmrE substrates lack the capacity to donate strong hydrogen bonds, reducing the geometric constraints for protein-substrate interactions. Prior MD simulations and NMR experiments suggested a dynamic interaction between TPP^+^ and the EmrE binding pocket(Shcherbakov et al., 2021; Vermaas et al., 2018), and we expect that many compounds transported by EmrE have some mobility within the binding pocket. In the present structural experiments, we observe this explicitly for methyl viologen, which we identified in different but overlapping positions in the two transporters in the asymmetric unit.

While our experiments indicate that altering sidechain configuration is important to accommodate diverse substrates, backbone conformational changes do not need to be invoked to explain polyspecificity. Indeed, we do not see perturbations in EmrE’s main chain structure in the six different EmrE crystal structures resolved here. In addition, the general correspondence of the structures of EmrE and Gdx-Clo indicates that same tertiary architecture can also accommodate substrates with guanidinyl headgroups and/or alkyl tails. These observations also jibe with observations from cryo-EM, which showed only minor differences in helix orientation and packing for the apo and TPP^+^-bound structures(Tate et al., 2003). Thus, the crystallized conformation can accommodate substrates from major classes, including quaternary ammoniums, quaternary phosphoniums, planar polyaromatics, and substituted guanidiniums, without substantial backbone rearrangement.

### Binding of benzalkonium^+^ and other substrates with alkyl chains

Because benzalkonium is especially relevant as a common household and hospital antiseptic to which the Qac proteins provide resistance, we sought to visualize how this quaternary ammonium compound might interact with EmrE. Although we were unable to generate diffracting crystals of EmrE_3_ in the presence of substrates with long alkyl tails, our current structure of EmrE_3_ with benzyltrimethylammonium bound (a chemical homologue of benzalkonium with a methyl group in place of the 10-12 carbon alkyl chain), combined with our previous Gdx-Clo structure, provides a strong indication of how benzalkonium or other detergent-like substrates might bind.

In Gdx-Clo, octylGdm^+^ binds such that its alkyl tail extends out of the aqueous binding pocket and into the membrane. In order to accommodate the alkyl tail, hydrophobic sidechains lining Gdx-Clo’s TM2 portal, including M39_Clo_ and F43_Clo_, adopted alternative rotamers(Kermani et al., 2020). Although all the substrates in the present EmrE_3_ structures were contained within the aqueous pocket, we similarly observe rotameric rearrangements of the TM2 sidechains in different structures, including Y40_EmrE_ and F44_EmrE_ (equivalent to Gdx-Clo’s M39_Clo_ and F43_Clo_) in the harmane and methyl viologen structures. These observations suggest that, as in Gdx-Clo, in EmrE the sidechain packing at the TM2 interface is malleable, and that movements of these residues may remodel the TM2 portal to permit binding of substrates with detergent-like alkyl chains.

Indeed, when the quaternary ammonium headgroup of benzalkonium is superposed onto the experimentally determined position of benzyltrimethylammonium in the EmrE_3_ binding pocket, the alkyl tail of benzalkonium extends towards the portal defined by the TM2 helices. Although the extended alkyl chain would clash with F44_B, EmrE_, positioning this sidechain in the “down” rotamer (analogous to that adopted by F43_B, Clo_ when octylguanidinium is bound) alleviates all clashes between the substrate and protein and provides unobstructed access for the alkyl tail to the membrane interior. Figure 7 shows a proposed model of benzalkonium binding to EmrE prepared by aligning the benzalkonium headgroup with benzyltrimethylammonium followed by energy minimization of the complex using MMTK(Hinsen, 2000).

**Figure 7.**
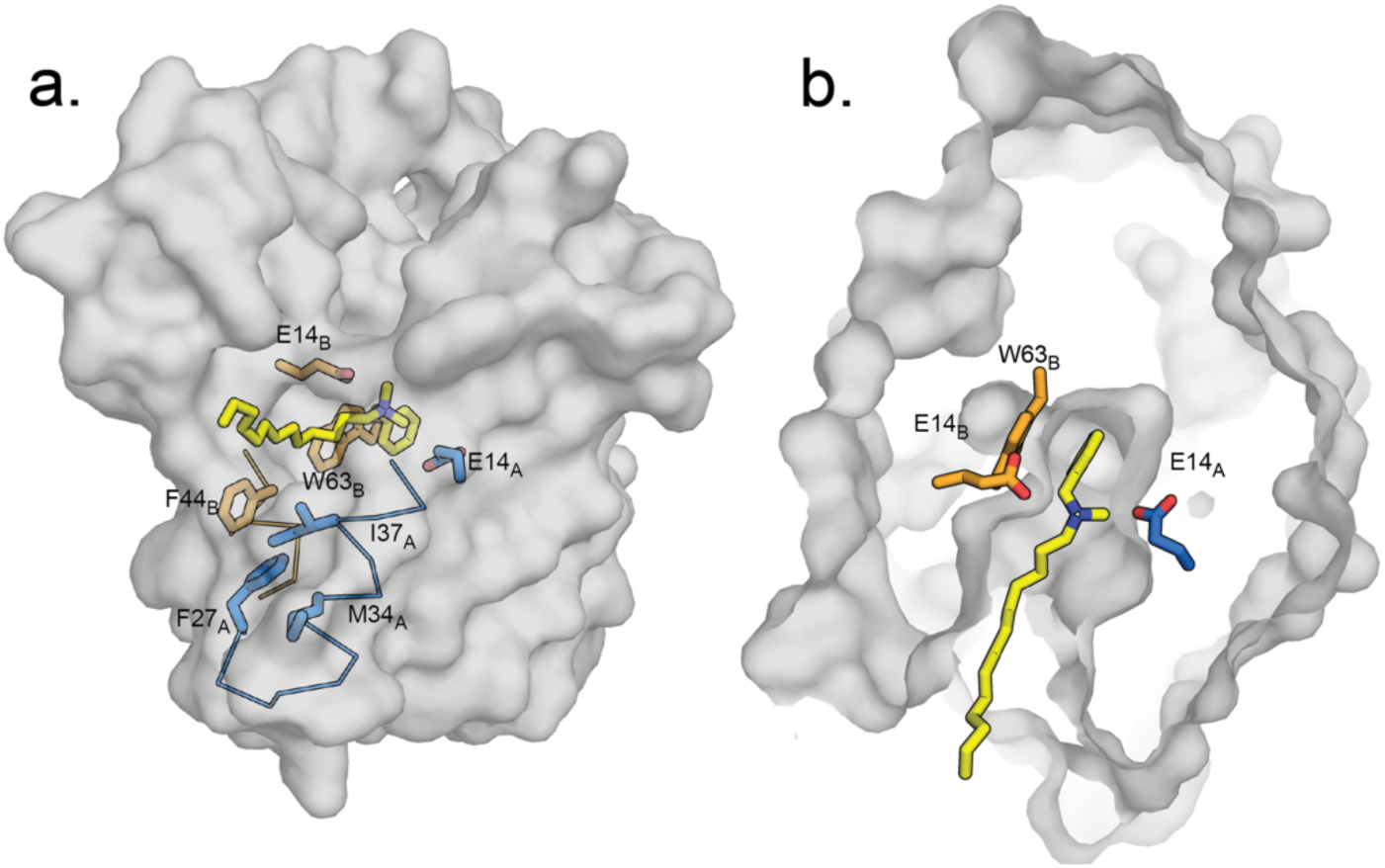
Hypothetical model of benzalkonium binding to EmrE. A. Benzalkonium is shown in yellow stick representation. Sidechains from the A and B subunits are colored in blue and orange as before. The mainchain for helices lining the TM2 portal is shown in ribbon format, with the portal-lining sidechains shown as sticks. B. Top-down view of binding site with benzalkonium. EmrE is sliced at the midpoint of the membrane. Experimental models of EmrE in complex with benzyltrimethylammonium (PDB:7T00) and Gdx-Clo in complex with octylguanidinium (PDB:6WK9) were used to construct this theoretical model; these experimental models are shown in Figure 7-Figure Supplement 1.

Thus, we propose that sidechain rearrangements along the membrane portal also contribute to substrate polyspecificity by allowing hydrophobic substituents to extend out of the substrate binding site and access the membrane interior. Similarly, we imagine that dipartite drugs transported by EmrE, such as propidium (a planar polyaromatic group linked to a tetraethyl ammonium) and dequalinium (two aromatic groups with a 10-carbon linker) may also utilize the portal for transport, with the protein-mediated transport of one moiety dragging its tethered lipophilic partner across the membrane.

## Conclusions

In summary, we have developed a multipurpose crystallization chaperone for SMR proteins and used this tool to resolve the first sidechain-resolution crystal structures of the bacterial SMR transporter, EmrE. In order to establish the structural basis of substrate polyspecificity, we resolved structures with five different substrates bound, including quaternary phosphoniums, planar aromatics, and a quaternary ammonium compound. We propose that, compared with more selective representatives of the SMR family, a relatively sparse hydrogen bond network among binding site residues in EmrE permits sidechain flexibility to conform to structurally diverse substrates.

## Materials and Methods

### Protein purification and crystallization

L10 monobody was purified from inclusion bodies exactly as described in detail previously(Kermani et al., 2020). pET15b plasmids bearing the EmrE_3_ coding sequence with an N-terminal hexahistidine tag and a thrombin cut site were transformed into *E. coli* C41 and grown overnight (15-18 hours) in Studier’s autoinduction media at 37°C. Pellets were resuspended in breaking buffer (50mM Tris-Cl pH 8.0, 100mM NaCl, 10mM tris(2-carboxyethyl)phosphine) (TCEP)) with 400μg DNase, 2mM MgCl_2_, 1mM PMSF, 1mg/mL lysozyme, 25μg pepstatin, and 500μg leupeptin. Resuspended pellets were lysed by sonication and extracted with 2% n-Decyl-β-D-Maltopyranoside (DM) (Anatrace) for 2 hours at room temperature. Extract was clarified by centrifugation (16000 rpm, 4°C, 45 minutes), and loaded onto TALON cobalt resin equilibrated with wash buffer (20mM tris-Cl pH 8.0, 100mM NaCl, 5 mM DM) supplemented with 5mM TCEP. Column was washed with wash buffer, and wash buffer supplemented with 10 mM imidazole before elution of EmrE_3_ with wash buffer supplemented with 400 mM imidazole. After exchange into wash buffer using PD-10 desalting columns (GE Healthcare) His tags were cleaved with thrombin (1 U/mg EmrE_3_) overnight at room temperature (21°C) prior to a final size exclusion purification step using a Superdex 200 column equilibrated with 10mM 2-[4-(2-hydroxyethyl)piperazin-1-yl]ethanesulfonic acid (HEPES) pH 7.5, 100mM NaCl, 4mM DM.

For functional measurements, protein was reconstituted by dialysis as previously described(Kermani et al., 2020). For SSM electrophysiology experiments, proteoliposomes were prepared with 20 mg EPL per ml, and a 1:20 protein:lipid mass ratio. Proteoliposomes were aliquoted and stored at -80° C until use. For crystallography of EmrE_3_, monobody L10 and EmrE_3_ were each concentrated to 10 mg/mL, and the L10 protein solution was supplemented with 4 mM DM. EmrE_3_ and L10 were combined in a 2.1:1 molar ratio and supplemented with lauryldimethylamine oxide (LDAO, final concentration of 6.6 mM). The protein solution was mixed with an equal volume of crystallization solution (0.3 μL in 96-well plates). Crystals formed after approximately 4 weeks, and were frozen in liquid nitrogen before data collection.

For crystallization with substrate, the EmrE_3_/monobody/LDAO solution was prepared as before, and substrate was added from a stock solution immediately before setting crystal trays (final concentrations of 1 mM for methyl viologen, 500 μM for harmane, 300 μM for benzyltrimethylammonium, 100 μM for TPP^+^, or 300 μM for MeTPP^+^). The low pH EmrE_3_ crystals grew in 200 mM NaCl, 100 mM sodium cacodylate, pH 5.2, 34% PEG 600. The substrate-bound EmrE_3_ crystals grew in 100 mM LiNO_3_ or 100 mM NH_4_SO_4_, 100 mM ADA, pH 6.5 or 100 mM HEPES, pH 7.1-7.3, and 30-35% PEG 600. Gdx-Clo protein and crystals were prepared exactly as described previously(Kermani et al., 2020). Crystals grew in 100 mM calcium acetate, 100 mM sodium acetate, pH 5.0, 40% PEG600.

### Structure determination

Crystallography data was collected at the Life Sciences Collaborative Access Team beamline 21-ID-D at the Advanced Photon Source, Argonne National Laboratory. Diffraction data were processed and scaled using Mosflm 7.3(Battye et al., 2011) or DIALS (Winter et al., 2018). Crystals diffracted anisotropically, and electron density maps were improved by anisotropic truncation of the unmerged data using the Staraniso webserver(Tickle, 2018) with a cutoff level of 1.2 -1.8 for the local *I/σ*. For the low pH EmrE_3_ dataset, phases were determined using molecular replacement with Phaser(McCoy et al., 2007), using the first three helices of Gdx-Clo and the L10 monobody structures (PDB:6WK8) as search models. Loop 3, helix 4, and the C-terminal loop were built into the experimental electron density using Coot(Emsley et al., 2010), with iterative rounds of refinement in Phenix(Liebschner et al., 2019) and Refmac(Murshudov et al., 2011). For the low pH Gdx-Clo structure, Gdx-Clo and the L10 monobody structures (PDB:6WK8) were used as molecular replacement search models. Models were validated using Molprobity(Williams et al., 2018) and by preparing composite omit maps in Phenix, omitting 5% of the model at a time(Terwilliger et al., 2008). The substrate-bound structures were phased using molecular replacement with monobody L10 and the A and B subunits of the initial EmrE_3_ model as the search models. Proteins typically crystallized in C121, although the methyl viologen-bound EmrE_3_ structure and the low pH Gdx-Clo crystallized in P1. For both, the unit cell contained two pseudosymmetric copies of the transporter-monobody complex.

### Microscale thermophoresis

Monobody L10 was labelled at a unique, introduced cysteine, A13C, with fluorescein maleimide. Binding to EmrE_3_ was measured using microscale thermophoresis (Nanotemper, Munich, Germany). For these experiments, labelled monobody was held constant at 2 μM, and the concentration of EmrE_3_ was varied from 30 nM - 100 μM. Buffer contained 100 mM NaCl, 10 mM HEPES, pH 7, 4 mM DM, and 50 μg/mL bovine serum albumin. Samples were incubated at least 30 minutes prior to measurement of binding interactions. Experiments were performed using three independent sample preparations and fit to a one site binding equilibrium with total L10 as the experimental variable:

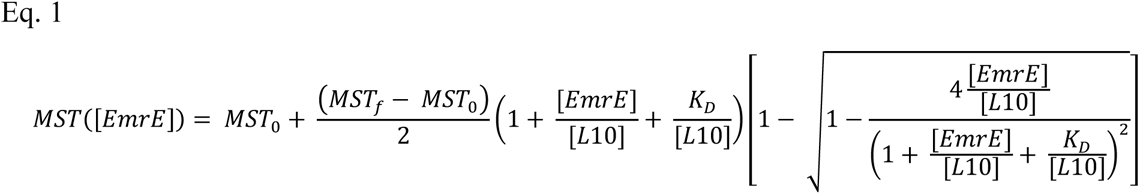

where MST([EmrE]) is the MST signal as a function of total EmrE added to a fixed concentration of labelled L10 monobody, and MST_0_ and MST_f_ are the arbitrary initial and final MST fluorescence signals.

### SSM electrophysiology

SSM electrophysiology was conducted using a SURFE^2^R N1 instrument (Nanion Technologies, Munich, Germany) according to published protocols(Bazzone and Barthmes, 2020; Bazzone et al., 2017). The sensor was alkylated and painted with lipid solution (7.5 µg/µl 1,2-diphytanoyl-sn-glycero-3-phosphocholine in n-decane), followed immediately by addition of recording buffer (100 mM KCl, 100 mM KPO_4_, pH 7.5). For measurements in the presence of monobody, buffers also contained 50 μg bovine serum albumin/mL. Proteoliposomes were applied to the sensor surface and centrifuged at 2500 x g for 30 minutes. Before experiments, sensors were checked for conductance and capacitance using SURFE^2^R software protocols. Sensors for which capacitance and conductance measurements were outside an acceptable range (10-40 nF capacitance, 1-5 nS conductance) were not used for experiments. Sensors were periodically rechecked for quality during the course of an experiment. When multiple measurements were performed on a single sensor, currents elicited by a reference compound were measured at the outset of the experiment and again after collecting data on test compounds. If currents differed by more than 10% between the first and last perfusions, this indicated that the proteoliposomes associated with the sensor had not remained stable over the course of the experiment, and data collected in this series was discarded. Between measurements, sensors were perfused with substrate-free solution for 2 s; observation of capacitive currents with opposite polarity indicated substrate efflux from the proteoliposomes and a return to the resting condition. For measurements of Michaelis-Menten kinetics, peak currents were used as a proxy for initial rate of transport(Bazzone et al., 2017). Titrations were performed on independent sensors; reported K_m_ values represent the mean of K_m_ values determined from fits to these independent titrations. In all cases, perfusion of protein-free liposomes with substrates did not evoke currents.

### NMR chemical shift prediction

The chemical shifts of the C*_α_* atoms of the NMR ensemble and the unliganded crystallography model were predicted using LARMOR^C^*^α^*(Frank et al., 2015) as implemented with PyShifts(Xie et al., 2020).

## Acknowledgements

We thank the Stockbridge lab for comments on the project and manuscript, and we are grateful to Aaron Frank (University of Michigan) for helpful conversations about chemical shift-based comparisons of the structures.

## Data availability

Atomic coordinates for the crystal structures have been deposited in the Protein Data Bank under accession numbers 7MH6 (EmrE_3_/L10), 7MGX (EmrE_3_/L10/methyl viologen), 7SVX (EmrE_3_/L10/harmane), 7SSU (EmrE_3_/L10/MeTPP^+^), 7SV9 (EmrE_3_/L10/TPP^+^), 7T00 (EmrE_3_/L10/benzyltrimethylammonium) and 7SZT (Gdx-Clo/L10).

## Author Contributions

Conceptualization, A.A.K., O.E.B., and R.B.S.; Methodology, A.A.K., O.E.B., A.K., S.K., and R.B.S.; Formal Analysis, A.A.K., O.E.B., and R.B.S.; Investigation, A.A.K., O.E.B., B.B.K., and A.K.; Writing – Original Draft, R.B.S.; Writing – Review and Editing, A.A.K., O.E.B., A.K., S.K., and R.B.S.; Visualization, A.A.K., O.E.B. and R.B.S.; Supervision, R.B.S.; Project Administration, R.B.S.; Funding Acquisition, S.K. and R.B.S.

---

**Figure 1 – Figure Supplement 1.**
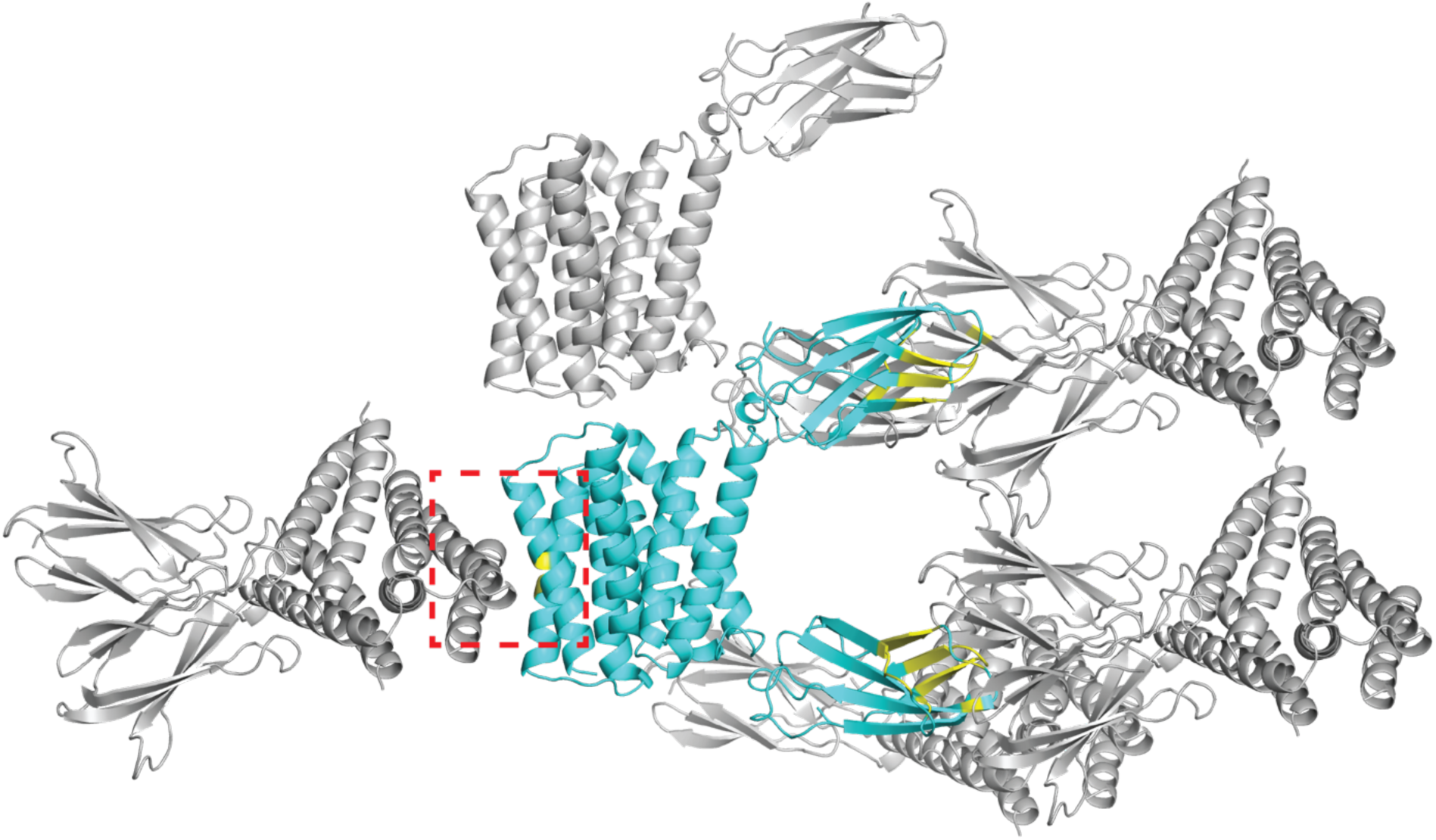
Crystal lattice for Gdx-Clo/L10 monobody complex (PDB: 6WK8). The asymmetric unit, composed of one Gdx-Clo dimer and two monobodies, is shown in cyan. Symmetry mates are shown in gray. Residues that contribute to an interface between the asymmetric unit and its symmetry mates are colored yellow. Five Gdx-Clo residues are in contact with a symmetry mate: TM4 residues V88_B_, L92_B_, T95_B_, F89_A_, L92_A_ (dashed red box).

**Figure 1 - Figure Supplement 2.**
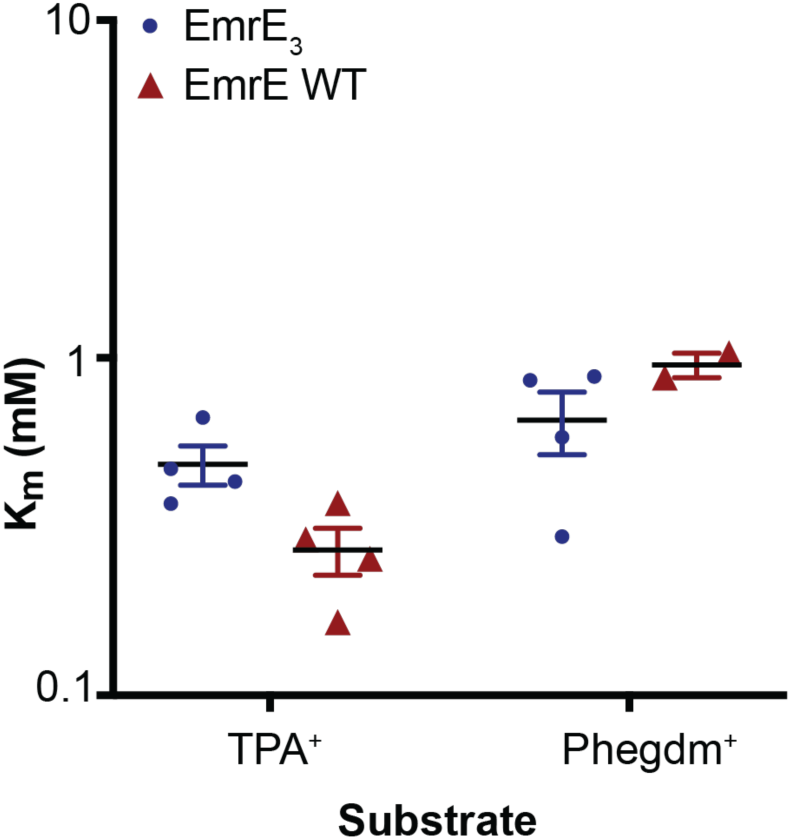
K_m_ values for TPA^+^ and PheGdm^+^ transport by EmrE_3_ (blue) and WT EmrE (red). Individual points are derived from Michaelis-Menten fits of titration experiments performed on a single sensor. Each K_m_ value was measured from a full titration series on an independently prepared sensor. Sensors are prepared from 2-3 independent biochemical purifications.

**Figure 1 – Figure Supplement 3.**
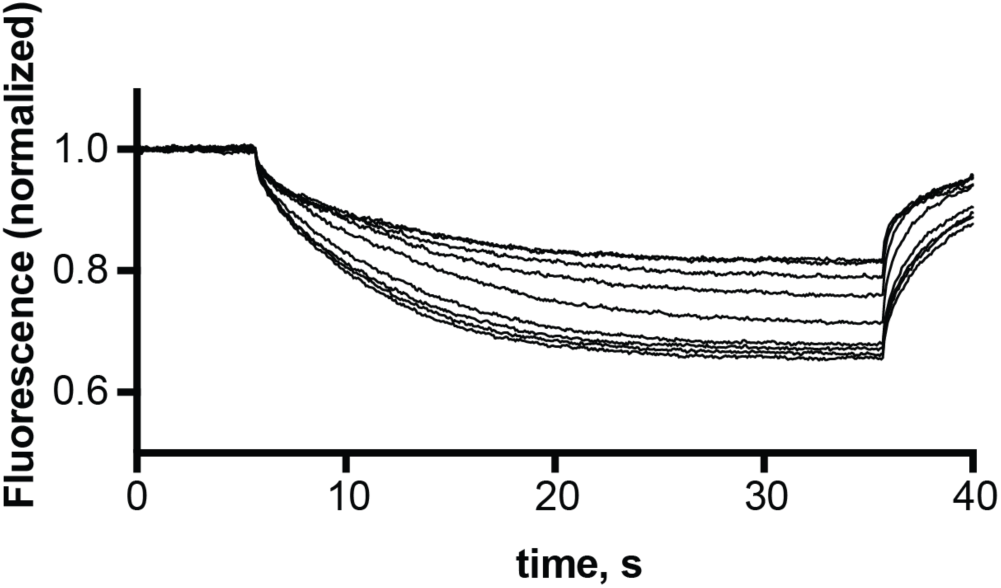
Representative microscale thermophoresis traces for monobody L10 in the presence of 30 nM – 10 μM EmrE_3_.

**Figure 2 — Figure Supplement 1.**
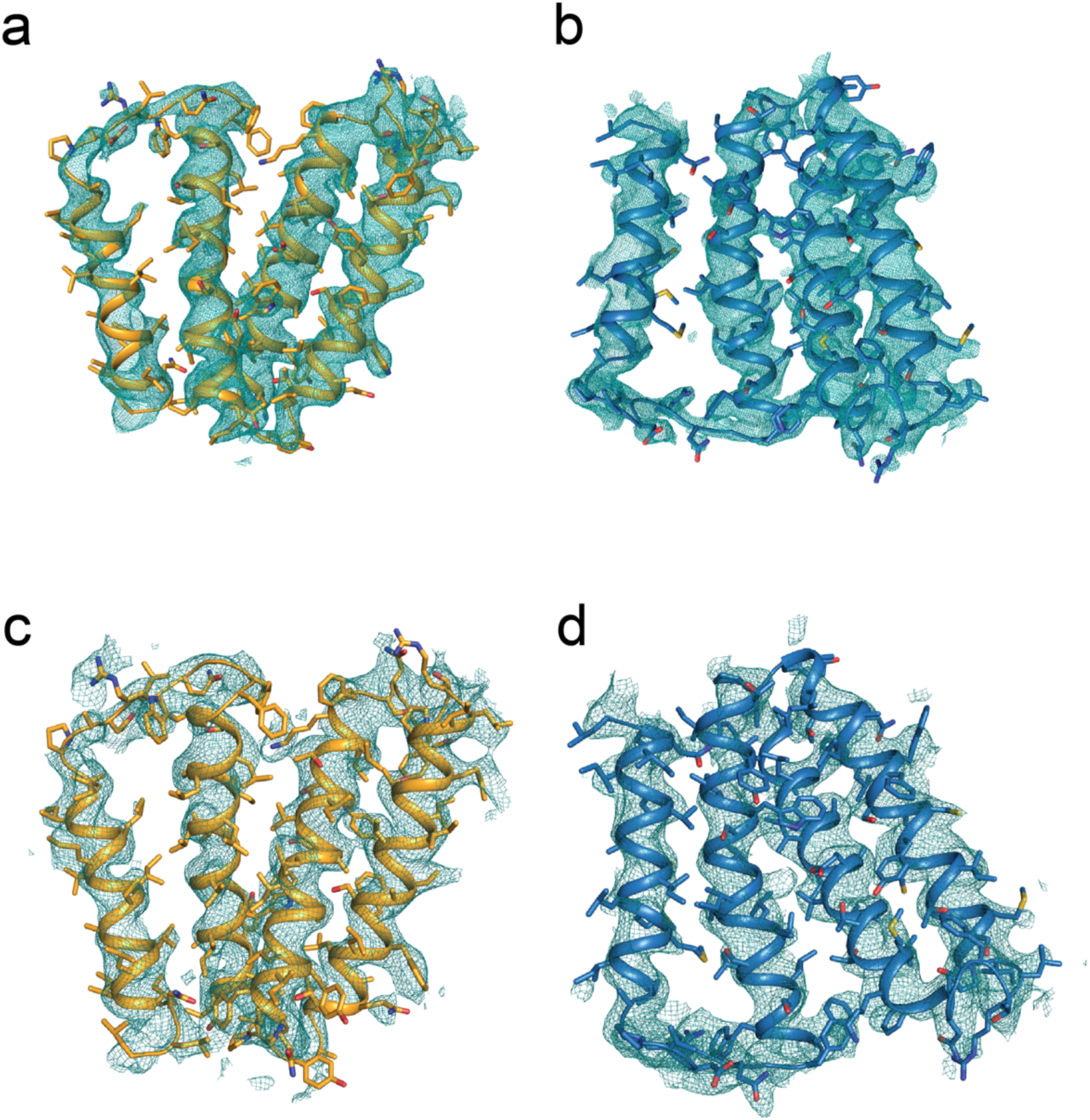
EmrE_3_ maps. Subunits colored as in main text, with subunit B in orange, and subunit A in blue. Panels A and B: 2F_o_-F_c_ maps for EmrE_3_, contoured at 1.2 *σ*. Panels C and D: 2F_o_-F_c_ composite omit maps for EmrE_3_, contoured at 1.0 *σ*, prepared by omitting 5% of the atoms in the model at a time.

**Figure 2 – Figure Supplement 2.**
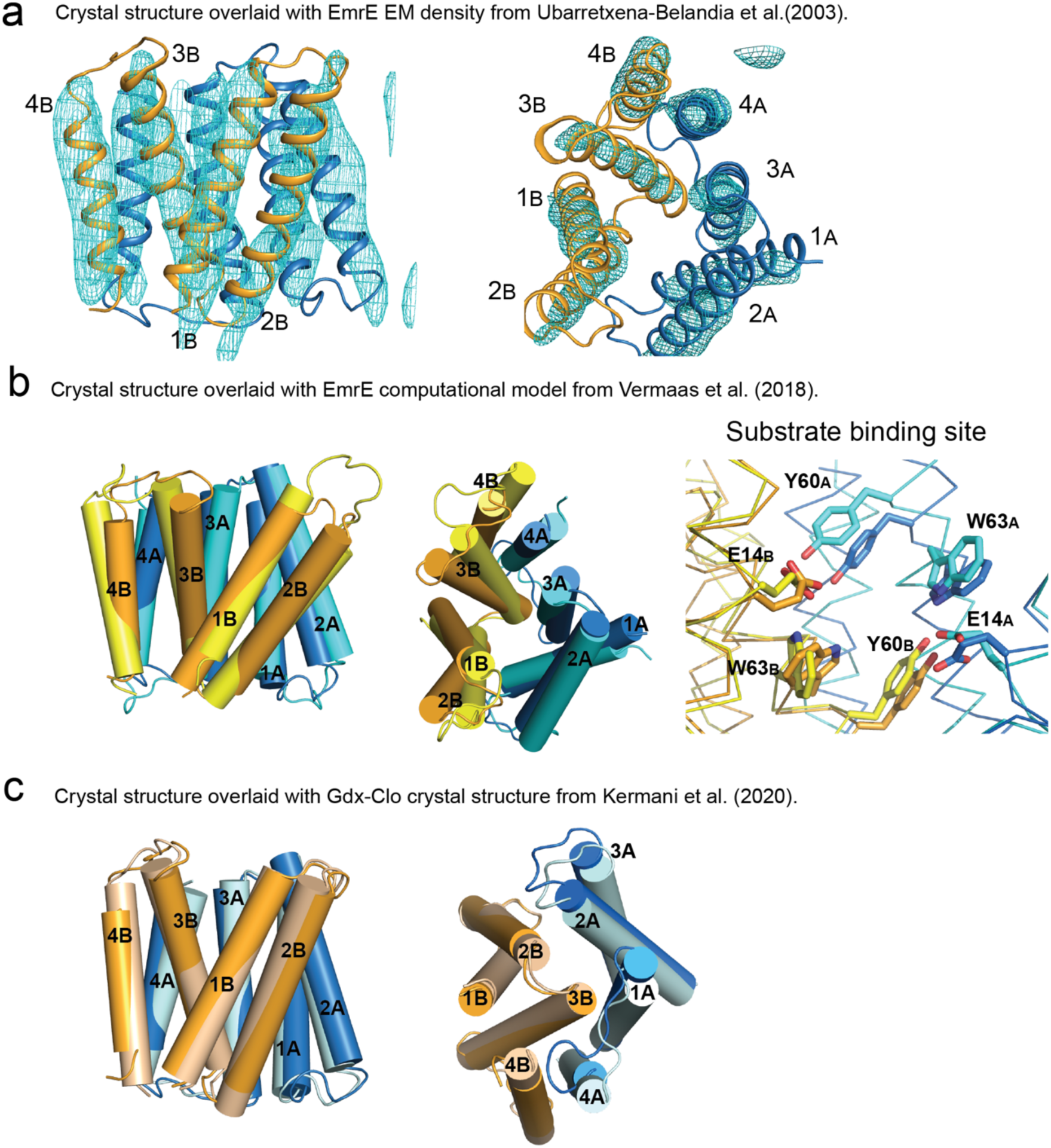
Structural comparison of EmrE_3_ crystal structure with electron microscopy maps, theoretical model, and Gdx-Clo. A. Crystal structure of EmrE_3_ (orange and blue cartoon) overlaid with experimental electron microscopy (EM) density (cyan mesh contoured at 1.5*σ*)(Ubarretxena-Belandia et al., 2003). B. Crystal structure of EmrE_3_ (orange and blue) compared to a computational model (yellow and cyan) constrained by EM data(Vermaas et al., 2018). C. Crystal structure of EmrE_3_ (orange and blue) compared to crystal structure of a homologue from the SMR family, Gdx-Clo (wheat and pale cyan)(Kermani et al., 2020). Models are aligned along the B subunit.

**Figure 2 – Figure Supplement 3.**
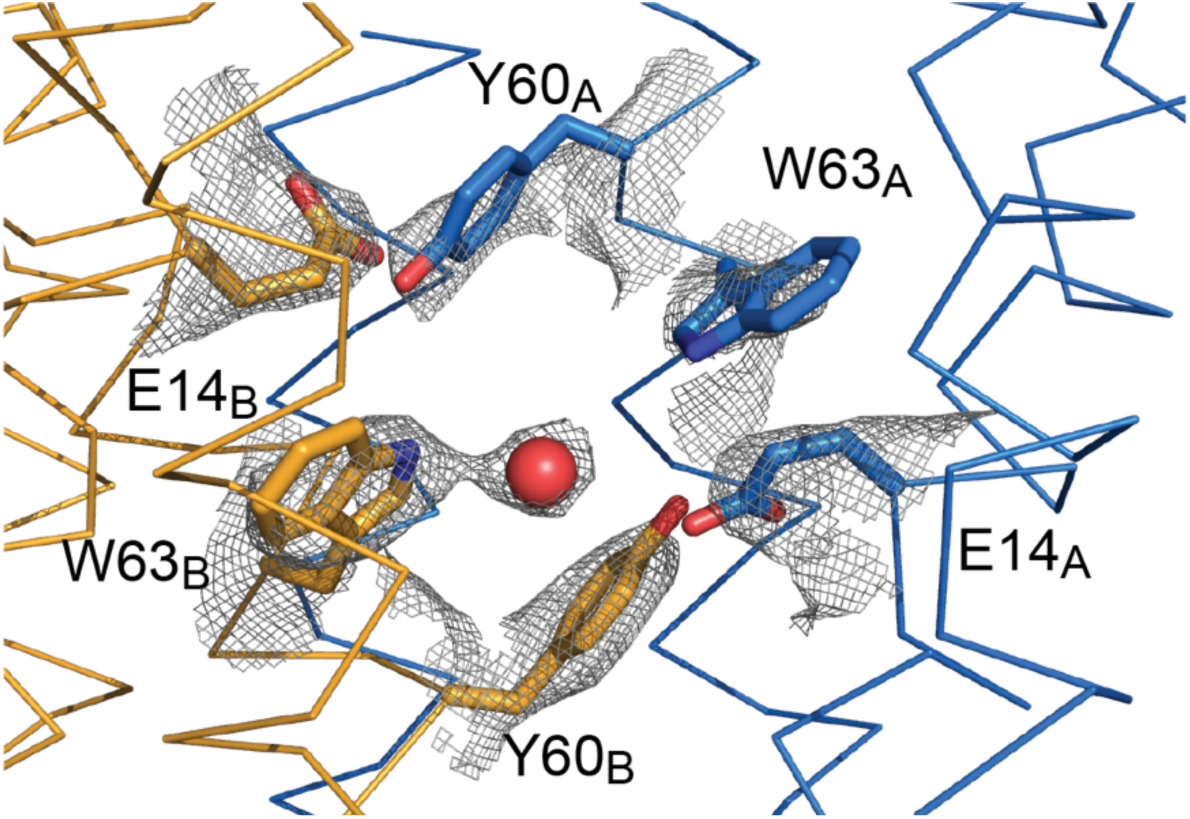
Sidechain density in the EmrE_3_ binding site. 2F_o_-F_c_ map around selected residues contoured at 1.5 *σ*. The red sphere represents a water molecule.

**Figure 3 – Figure Supplement 1.**
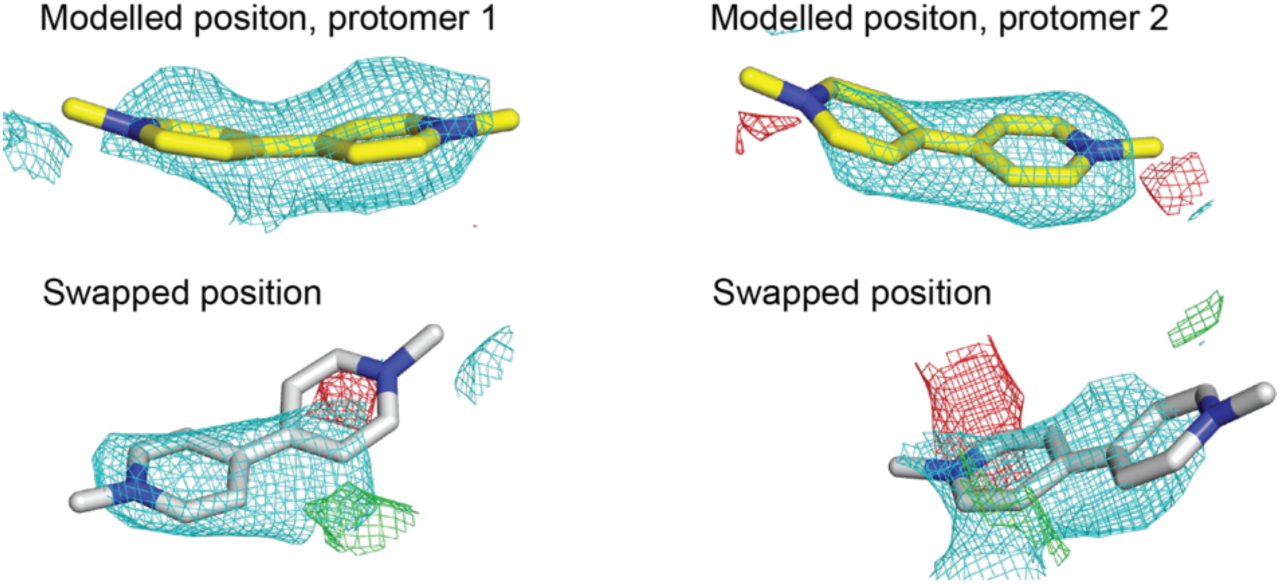
Electron density maps of methyl viologen in different EmrE_3_ protomers in the asymmetric unit. Yellow stick representations (top panels) show the modelled position of methyl viologen in each protomer, with the final refined maps shown as mesh. White stick representations show the methyl viologen position swapped between the two protomers. Maps show a subsequent re-refinement with the substrates in the swapped positions. For all panels, 2F_o_-F_c_ density (cyan) contoured at 1.2*σ*and F_o_-F_c_ density (green or red) contoured at 2.5 *σ*.

**Figure 3-Figure Supplement 2.**
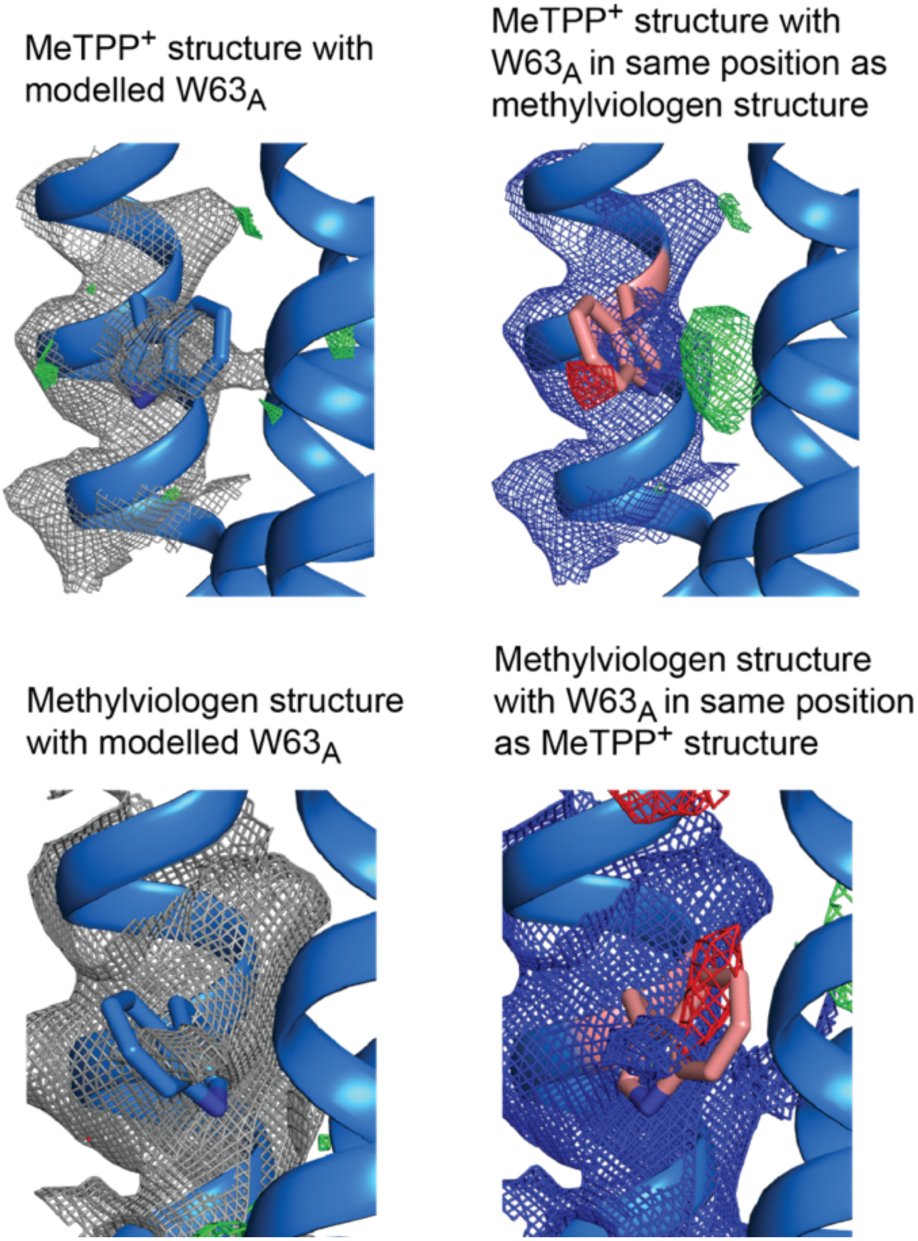
Electron density maps for W63_A_ modelled in different positions. Top panels: 2F_o_-F_c_ density contoured at 1.8*σ* and F_o_-F_c_ density contoured at 3 *σ*. Bottom panels: 2F_o_-F_c_ density contoured at 1.2 *σ* and F_o_-F_c_ density contoured at 2.5 *σ*.

**Figure 3-Figure Supplement 3.**
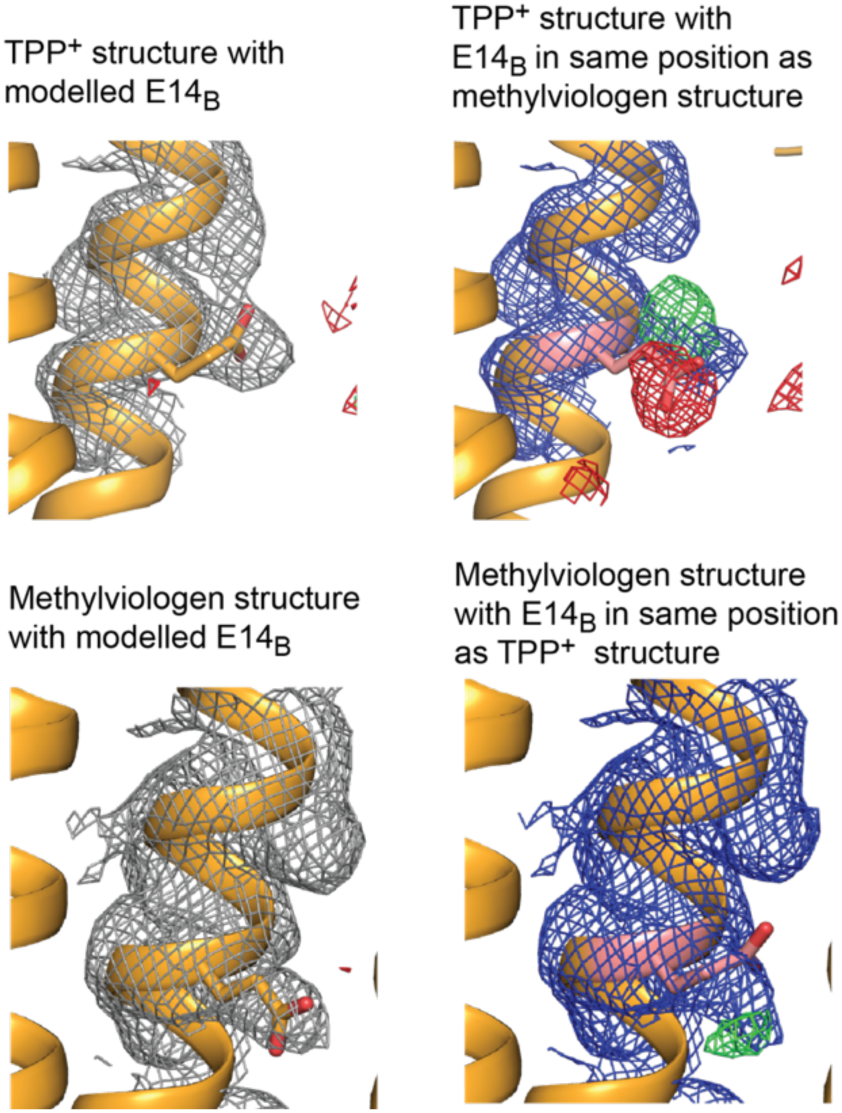
Electron density maps for E14_B_ modelled in different positions. Top panels: 2F_o_-F_c_ density contoured at 1.8*σ* and F_o_-F_c_ density contoured at 3 *σ*. Bottom panels: 2F_o_-F_c_ density contoured at 1.2 *σ* and F_o_-F_c_ density contoured at 2.5 *σ*.

**Figure 4- Figure Supplement 1.**
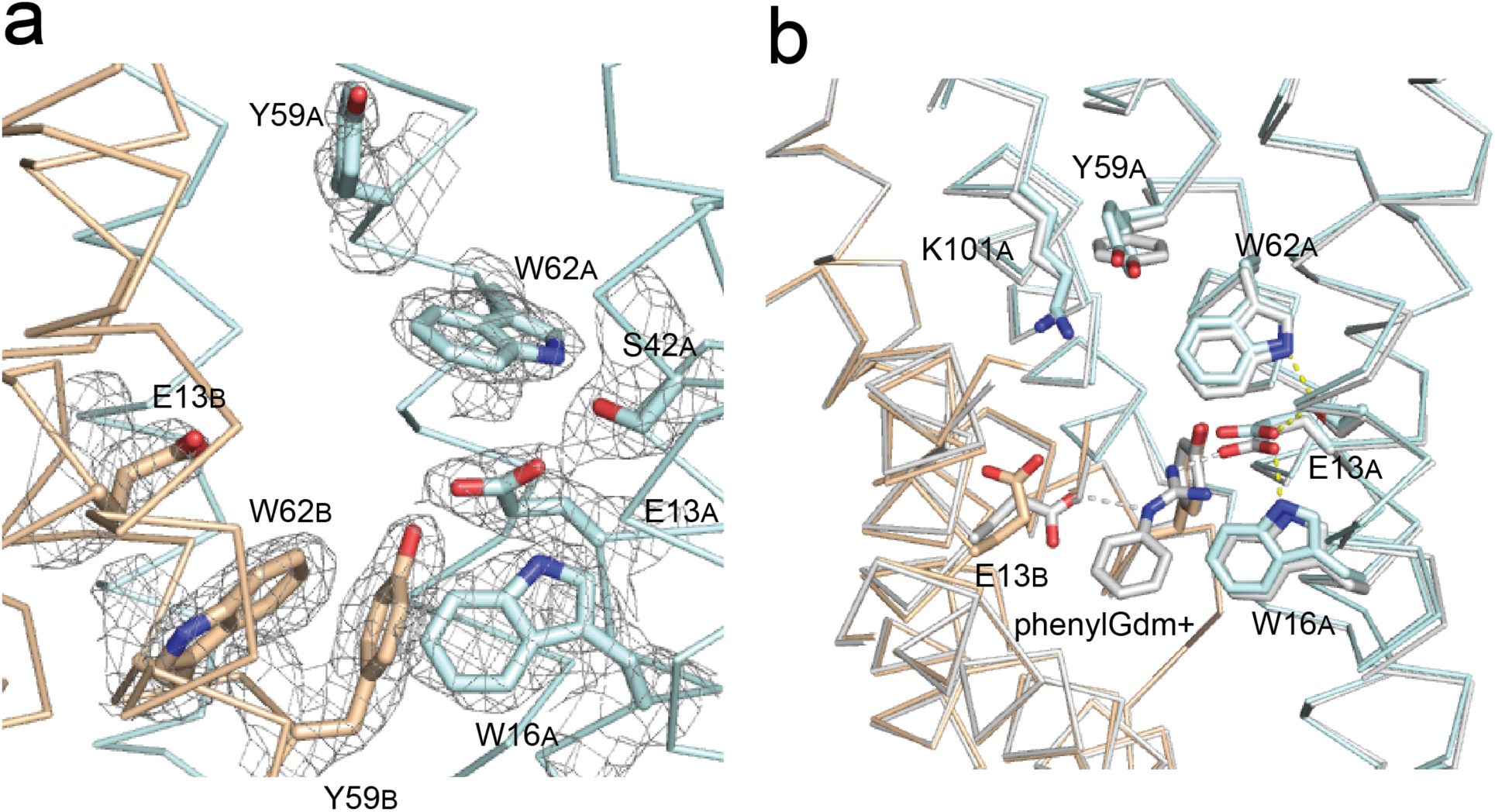
Gdx-Clo and EmrE substrate binding sites. A. 2F_o_-F_c_ map shown around selected residues in the Gdx-Clo substrate binding site (pH 5.2) contoured at 1.5*σ*. B. Alignment of Gdx-Clo structures. The present pH 5.2 structure is shown in wheat and cyan with putative H-bond interactions shown as yellow dashed lines. The structure with phenylGdm^+^ bound (PDB:6WK8) is shown in light gray with putative H-bonds between the substrate and the E13 residues shown as gray dashed lines.

**Figure 6- Figure Supplement 1.**
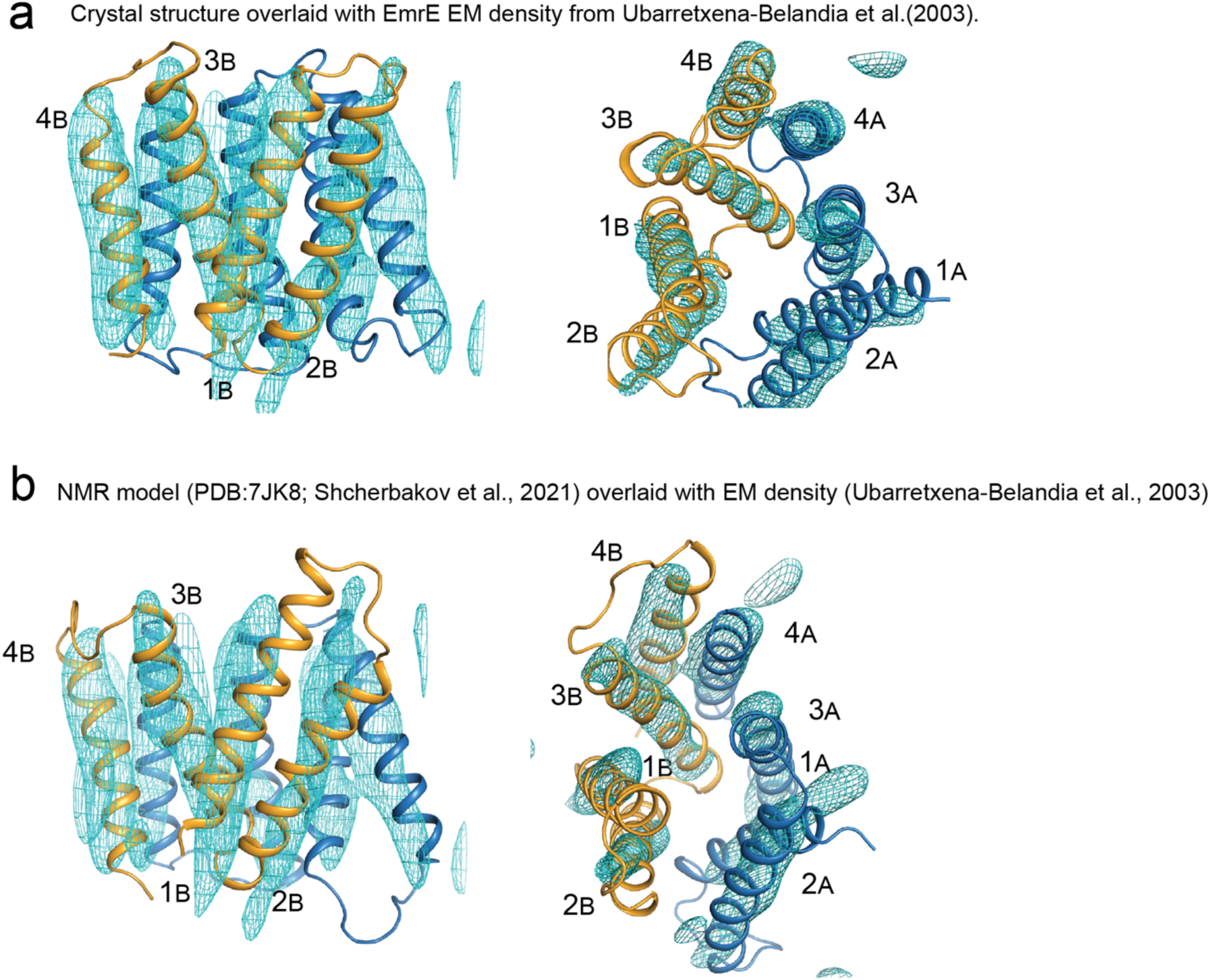
A. Crystal structure of EmrE_3_ (orange and blue cartoon) overlaid with experimental electron microscopy density (cyan mesh contoured at 1.5*σ*)(Ubarretxena-Belandia et al., 2003). (Panel repeated from Figure 2-Supplement 2 to aid visual comparison). B. NMR model of EmrE S64V (orange and blue cartoon)(Shcherbakov et al., 2021) overlaid with experimental electron microscopy density shown in panel A (cyan mesh contoured at 1.5*σ*)(Ubarretxena-Belandia et al., 2003).

**Figure 6- Figure supplement 2.**
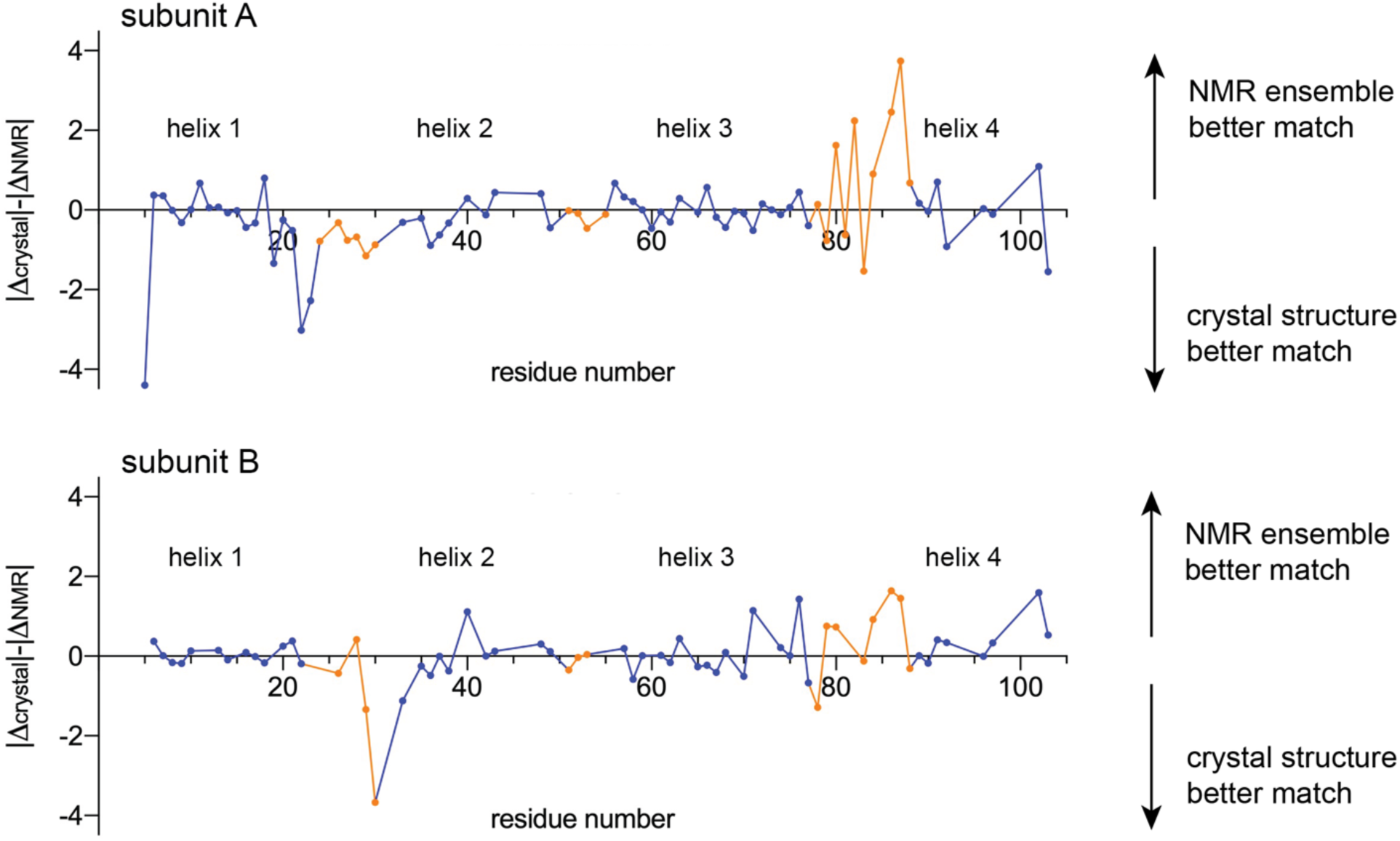
Comparison of experimental chemical shifts for EmrE (BMRB accession number 50411) with chemical shifts predicted from the crystallography model and NMR ensemble using LARMOR^C^*^α^*(Frank et al., 2015). Residue number is plotted along the x-axis. The y-axis compares the relative difference between the experimental chemical shifts and the predicted chemical shifts for the crystallography and NMR models. For each C*_α_*position, the difference between the predicted and experimental chemical shifts was calculated (*δ*_predicted, NMR model_-*δ*_experimental_ = *Δ*_NMR_ and *δ*_predicted, crystallography model_-*δ*_experimental_ = *Δ*_crystal_), and their relative magnitude compared (|*Δ*_crystal_|-|*Δ*_NMR_|). Values above the origin line indicate that the experimental chemical shifts are in better agreement with the predicted chemical shifts for the NMR ensemble; values below the origin line indicate that the experimental chemical shifts are in better agreement with the predicted chemical shifts for the crystallography model. Residues in TM helices are shown as blue points, and residues in loop regions are shown as orange points. Residues that were not assigned in the NMR dataset, or that are mutated in either the NMR or crystal structures (E25, W31, V34, S64) are absent from this plot.

**Figure 7 – Figure Supplement 1.**
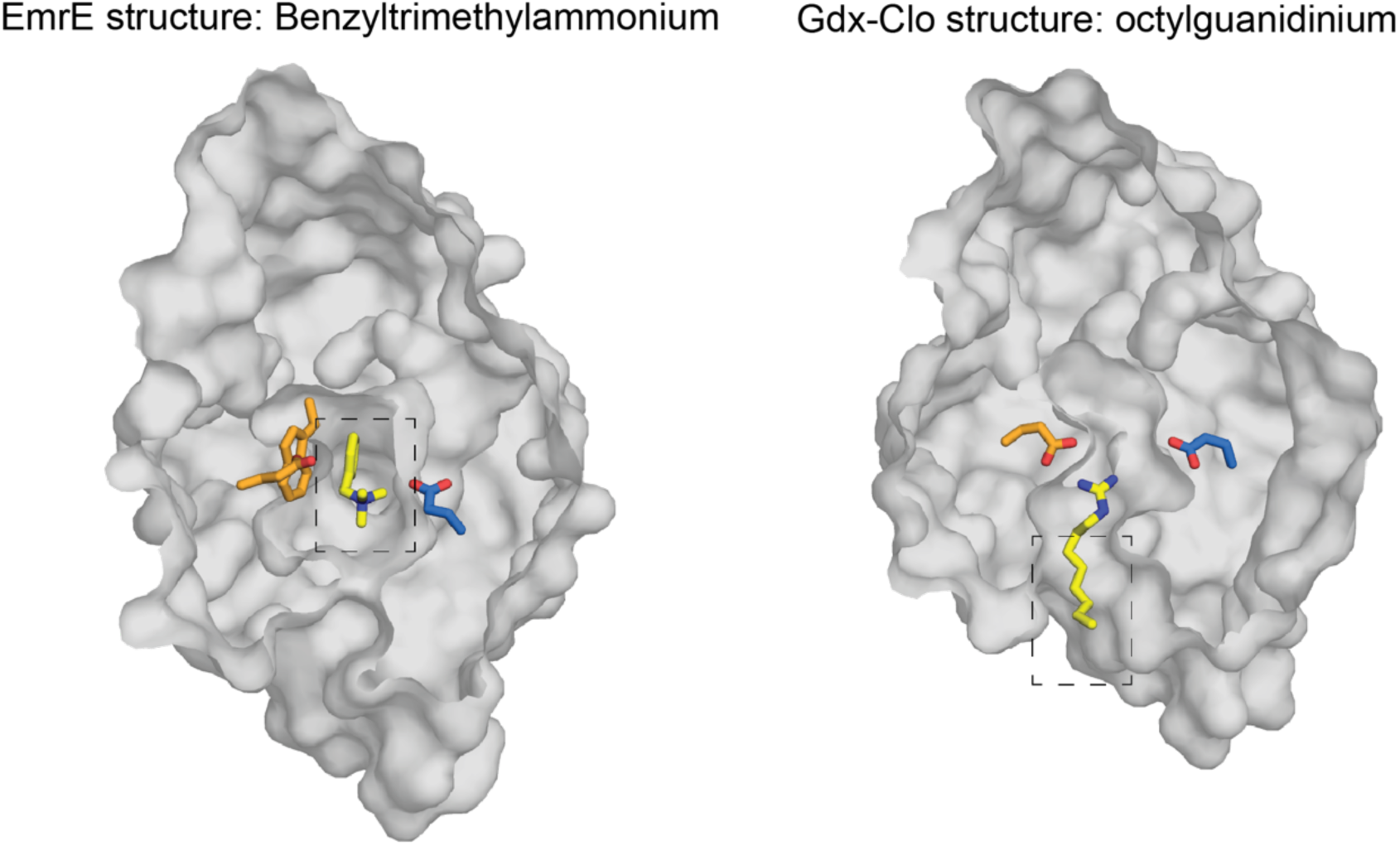
Top down structures of EmrE in complex with benzyltrimethylammonium (PDB:7T00; model for benzalkonium headgroup binding) and Gdx-Clo in complex with octylguanidinium (PDB:6WK9; model for alkyl tail positioning). Structures are sliced at the midpoint of the membrane, as in Figure 7. Dashed boxes indicate the headgroup and alkyl group positions used to prepare the hypothetical model of benzalkonium binding.

